# The zinc transporter Slc30a1 in macrophages plays a protective role against *Salmonella* infection

**DOI:** 10.1101/2023.06.06.543958

**Authors:** Pinanong Na-Phatthalung, Shumin Sun, Enjun Xie, Jia Wang, Junxia Min, Fudi Wang

## Abstract

The zinc transporter Slc30a1 plays an essential role in maintaining cellular zinc homeostasis; however, its functional role in macrophages remains largely unknown. Here, we systematically examined the expression and function of Slc30a1 in macrophages upon *Salmonella* infection in both Slc30a1 reporter mice and in macrophage-specific *Slc30a1* knockout (*Slc30a1^fl/fl^LysM^Cre^*) mice. We found that *Slc30a1^fl/fl^LysM^Cre^*mice have an increased susceptibility to *Salmonella* infection compared to control littermates. Mechanistically, we found that loss of Slc30a1 in macrophages reduced their bactericidal activity via reduced iNOS and NO production due to intracellular zinc accumulation. In addition, we observed significantly increased expression of *Mt1* (metallothionein 1) in *Salmonella*-infected *Slc30a1*-deficient macrophages, suggesting that Mt1 may serve as a compensatory zinc reservoir. Interestingly, macrophages lacking both *Mt1* and *Slc30a1* expression (*Slc30a1^fl/fl^LysM^Cre^; Mt1^-/-^*) had increased cell death upon *Salmonella* infection due to excess zinc-induced oxidative stress. Taken together, our results show that Slc30a1 in macrophages can protect against *Salmonella* infection, providing mechanistic insights into the role of Slc30a1-mediated zinc homeostasis in macrophages in response to infectious disease.

## Introduction

*Salmonella*, the gram-negative bacterium that causes Salmonellosis, a well-characterized and relatively common foodborne disease. According to the 2015 Global Burden of Disease Study, *Salmonella* infection is one of the leading causes of diarrhea-related death, particularly among children under 5 years of age *(GBD, 2017)*. In severe cases with an invasive infection of the bloodstream, fatality can reach as high as 25% *(Feasey et al., 2012)*. Based on epidemiological reports, zinc has been linked to the incidence of *Salmonella* infection *(GBD, 2017)*. As an essential micronutrient, zinc is involved in a wide range of fundamental biological processes, including modulating the immune system’s response to invading pathogens *(Wessels et al., 2021)*. However, whether zinc plays a protective role in response to *Salmonella* infection remains unclear.

In mammalian cells, zinc homeostasis is tightly regulated by two families of zinc transporters, namely solute carrier family 30 (Slc30a, also known as ZnTs) and solute carrier family 39 (Slc39a, also known as ZIPs), which export and import cellular zinc, respectively. Zinc imbalance caused by dysfunction of zinc transporters contributes to impairment of several important functions, such as hematopoiesis and macrophage survival *(He et al., 2023; Gao et al., 2017)*. Slc30a1 (ZnT1) is ubiquitously expressed in mammalian cells and is localized primarily at the plasma membrane *(Nishito and Kambe, 2019; Golan et al., 2015; Guo et al., 2010)*, where it helps maintain cellular zinc homeostasis by exporting cytoplasmic zinc to the extracellular space *(Shusterman et al., 2017, Kambe et al., 2015, Kimura and Kambe, 2016)*. In mice, loss of Slc30a1 causes early embryonic death *(Andrews et al., 2004)*, suggesting that Slc30a1 is essential for early development and survival. Recently, Slc30a1 expression was reported to be induced by a wide range of pathogens, including bacteria *(Botella et al., 2011; Stocks et al., 2021)*, fungi *(Rossi et al., 2021)*, and viruses *(Moskovskich et al., 2019)*.

Macrophages serve as the front line in the innate immune system by detecting and eliminating foreign pathogens. When *Salmonella* invades the epithelium, the bacteria are rapidly engulfed and killed by resident macrophages. This rapid engulfment and killing of *Salmonella* by macrophages can have potentially important consequences such as bacterial clearance, cytokine production, and the recruitment of polymorphonuclear phagocytes *(Mastroeni et al., 2009)*. Despite evidence suggesting a link between zinc regulators and these macrophage functions *(Gao et al., 2018)*, whether and how Slc30a1 regulates macrophage function during infection remains poorly understood.

Here, we performed RNA-seq analysis and found that *Slc30a1* expression is upregulated in *Salmonella*-infected macrophages. To study the role of Slc30a1 in macrophages upon *Salmonella* infection, we then generated macrophage-specific *Slc30a1* knockout mice (*Slc30a1^fl/fl^LysM^Cre^*) and found that these mice are more susceptible to *Salmonella* infection compared to control littermates. In addition, we found that *Slc30a1*-deficient macrophages have impaired bactericidal capacity due to reduced production of iNOS (inducible nitric oxide synthase) and NO. Based on our finding that *Mt1* (which encodes the metalloprotein metallothionein 1) is significantly upregulated in *Slc30a1^fl/fl^LysM^Cre^*mice upon *Salmonella* infection, we then generated macrophage-specific *Slc30a1* and *Mt1* double-knockout mice (DKO) by crossing *Slc30a1^fl/fl^LysM^Cre^* mice with global *Mt1*-deficient (*Mt1^-/-^*) mice. To our surprise, we found that *Mt1* promotes cell death, but has no effect on either cellular zinc levels or antimicrobial activity compared to *Slc30a1*-deficient macrophages upon *Salmonella* infection. These results provide novel mechanistic insights into the protective role of Slc30a1 and its ability to maintain zinc homeostasis in macrophages in response to pathogen infection, suggesting that targeting Slc30a1 may provide new treatment strategies for managing *Salmonella* infection.

## Results

### *Slc30a1* expression is induced in mouse macrophages upon *Salmonella* infection

Our first step was to perform RNA sequencing (RNA-seq) on bone marrow-derived macrophages (BMDMs) that were isolated from wild-type C57BL/6 mice and then exposed to *Salmonella* with a multiplicity of infection (MOI) of 1 for two hours (Figure 1A), the approximate time that has been closely correlated with the initial replication of *Salmonella* in macrophages *(Helaine et al., 2010)*. Our analysis revealed a total of 1074 differentially expressed genes (DEGs), defined as an absolute log_2_ fold change >1 and an adjusted *P*-value of <0.05 compared to control-treated BMDMs (Figure 1B). Among these 1074 genes, significantly downregulated genes were enriched in anti-inflammatory pathways, including *Gpx1*, *Smad7*, and *Tgfb3*, and significantly upregulated genes were enriched in infectious and inflammatory pathways, including *Tnf*, *Il6*, *Il1β*, *Ptgs2*, and *Nos2*, suggesting a potent antimicrobial response in infected macrophages (Figure 1C); as shown in a heatmap, a large number of genes were upregulated following *Salmonella* infection (Figure 1D). Gene ontology (GO) analysis of the DEGs revealed that the top 50 GO terms generally contributed to proinflammatory signaling, inflammatory cytokines, and responses to molecules of bacterial origin (Figure supplement 1 and Table supplement 1), while Kyoto Encyclopedia of Genes and Genomes (KEGG) pathway enrichment analysis of these DEGs yielded 54 pathways (*P*<0.05) involved largely in infectious diseases (Figure 1E and Table supplement 2).

**Figure 1.**
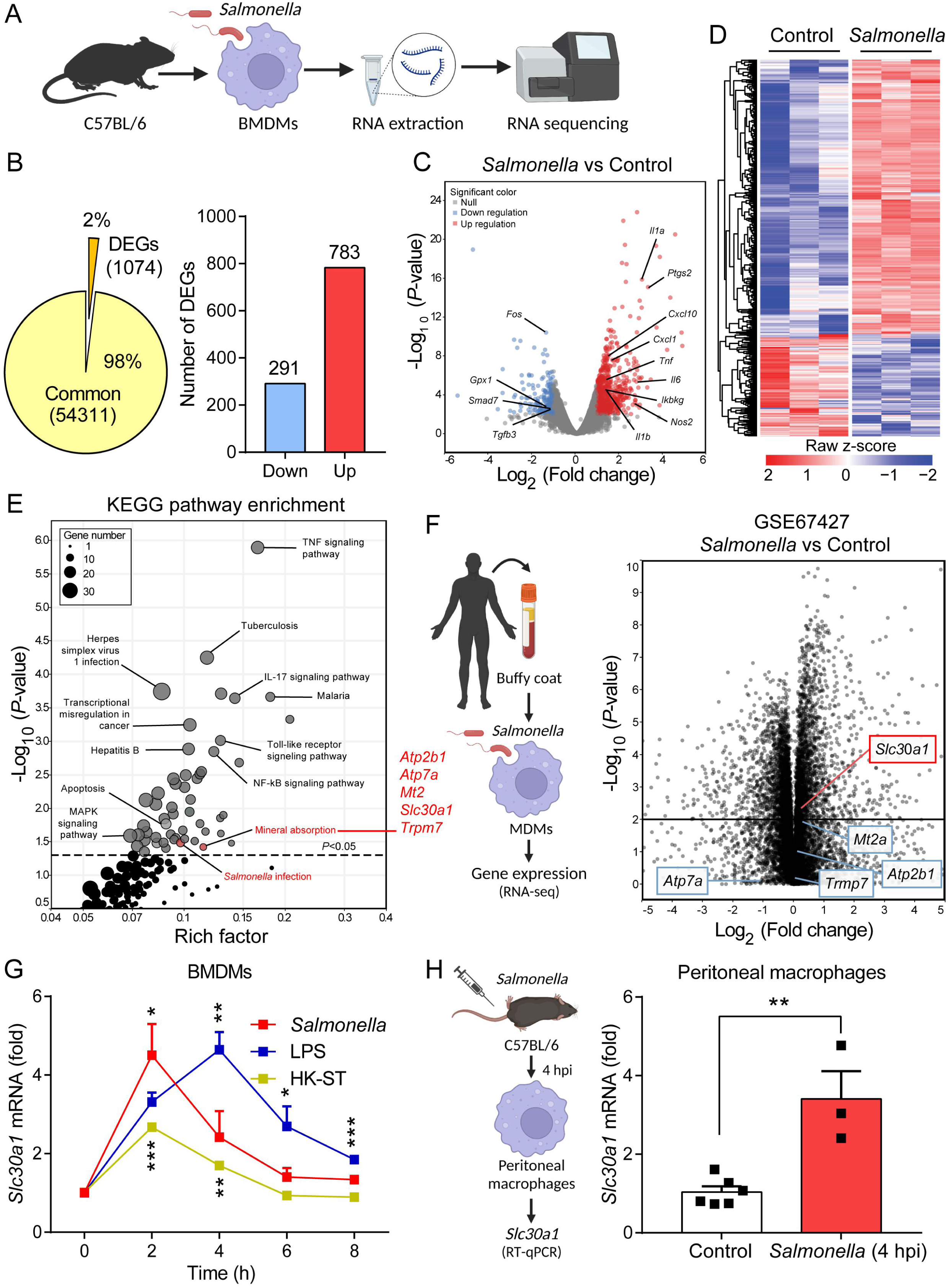
*Slc30a1* expression is upregulated in *Salmonella*-infected macrophages. **(A)** Schematic diagram illustrating the experimental design for measuring expression using RNA-seq in wild-type (WT) bone marrow-derived macrophages (BMDMs) 2 h after *Salmonella* infection (MOI = 1) (*n* = 3). (**B**) Pie chart (left) summarizing the percentage of unchanged genes and differentially expressed genes (DEGs), defined as *P*<0.05 and an absolute log_2_ fold change >1; shown at the right is the total number of downregulated and upregulated genes. (**C**) Volcano plot displaying the fold change in gene expression and corresponding *P*-values for *Salmonella*-infected cells vs. control (uninfected) cells. (**D**) Heatmap showing the pattern of DEGs in BMDMs measured between *Salmonella*-infected cells and control cells. (**E**) KEGG enrichment pathways (*P*<0.05) of DEGs between *Salmonella*-infected cells and control cells. (**F**) Schematic diagram (left) illustrating the use of RNA-seq analysis on human monocyte-derived macrophages (MDMs) infected with *Salmonella* for 4 h and volcano plot (right) of DEGs in *Salmonella*-infected MDMs based on a previously published dataset (GSE67427); the *Atp2b1*, *Atp7a*, *Mt2*, *Slc30a1*, and *Trpm7* genes are indicated. (**G**) Time course of *Slc30a1* mRNA measured using RT-qPCR in mouse BMDMs infected with *Salmonella* (MOI = 1), stimulated with lipopolysaccharide (LPS, 100 ng/ml), or infected with heat-killed *Salmonella* (HK-ST, MOI = 1) (*n* = 3). (**H**) Schematic diagram (left) illustrating the strategy for measuring *Slc30a1* mRNA in peritoneal macrophages isolated from WT mice 4 h after infection with 10^5^ CFU *Salmonella* (4 hpi) or uninfected control mice. Shown at the right is the summary of *Slc30a1* mRNA measured in peritoneal macrophages (each symbol represents a pooled sample from 8-12 mice). Data in G and H are presented as mean ± SEM. *P* values in G and H were determined using 2-tailed unpaired Student’s *t-*test. \**P*<0.05, \*\**P*<0.01 and \*\*\**P*<0.001.

Using KEGG pathway analysis, we identified mineral absorption as a predominant pathway, with upregulation of the *Atp2b1*, *Atp7a*, *Mt2*, *Slc30a1*, and *Trpm7* genes. To narrow down the list of genes, we analyzed publicly available bulk RNA-seq data (GSE67427) obtained from human monocyte-derived macrophages that were infected for 4 h with *Salmonella typhimurium* and found that the DEGs in our mouse RNA-seq dataset were also differentially regulated in human macrophages (Figure 1F). Importantly, *Slc30a1* was ranked as the most significantly upregulated genes and was therefore selected for further study. To validate the RNA-seq data, we then used RT-qPCR to measure *Slc30a1* mRNA levels in BMDMs at various time points following exposure to *Salmonella*, lipopolysaccharide (LPS), or heat-killed *Salmonella typhimurium* (HK-ST). We found that *Slc30a1* expression peaked at 2 and 4 h in the *Salmonella*- and LPS-treated cells, respectively, while HK-ST produced a significantly smaller response that also peaked at 2 h (Figure 1G). To validate this *in vitro* finding *in vivo*, we gave C57BL/6 mice an intraperitoneal (i.p.) injection of a sublethal dose of *Salmonella* (1×10^5^ CFU per mouse); at 4 hours post-infection (4 hpi), *Slc30a1* expression was significantly higher in peritoneal macrophages compared to uninfected mice (Figure 1H). Together, these data show that *Slc30a1* is upregulated in macrophages during *Salmonella* infection.

### Macrophages in *Salmonella*-infected Slc30a1 reporter mice have increased levels of Slc30a1 protein

To detect the levels of Slc30a1 protein in macrophages following *Salmonella* infection, we generated an Slc30a1 reporter mouse line expressing 3xFlag-2A-EGFP-2A-CreERT2-Wpre-pA under the control of the *Slc30a1* promoter (Figure 2A, B) and studied heterozygous offspring mice (*Slc30a1^flag-EGFP/+^*). BMDMs were then isolated from *Slc30a1^flag-EGFP/+^* mice and either infected with *Salmonella* (MOI = 1) or treated with ZnSO_4_ (40 µM) for 4 h, followed by western blot analysis using an anti-flag antibody. Compared to untreated WT and untreated *Slc30a1^flag-EGFP/+^* BMDMs, both the *Salmonella*-infected and ZnSO_4_-treated *Slc30a1^flag-EGFP/+^* BMDMs had significantly higher levels of the reporter protein (Figure 2C). Given that Slc30a1 localizes primarily to the plasma membrane *(Nishito and Kambe, 2019; Golan et al., 2015; Guo et al., 2010)*, we examined the localization of Slc30a1 in *Salmonella*-infected and ZnSO_4_-treated cells using confocal microscopy. As expected, we found that ZnSO_4_ increased the levels of the Slc30a1-flag reporter, primarily in the plasma membrane (Figure 2D). Interestingly, however, the Slc30a1-flag reporter was present both in the cytosol and in the plasma membrane in *Salmonella*-infected cells (Figure 2D), suggesting that Slc30a1 may contribute to both zinc efflux and intracellular zinc movement in response to *Salmonella* infection.

**Figure 2.**
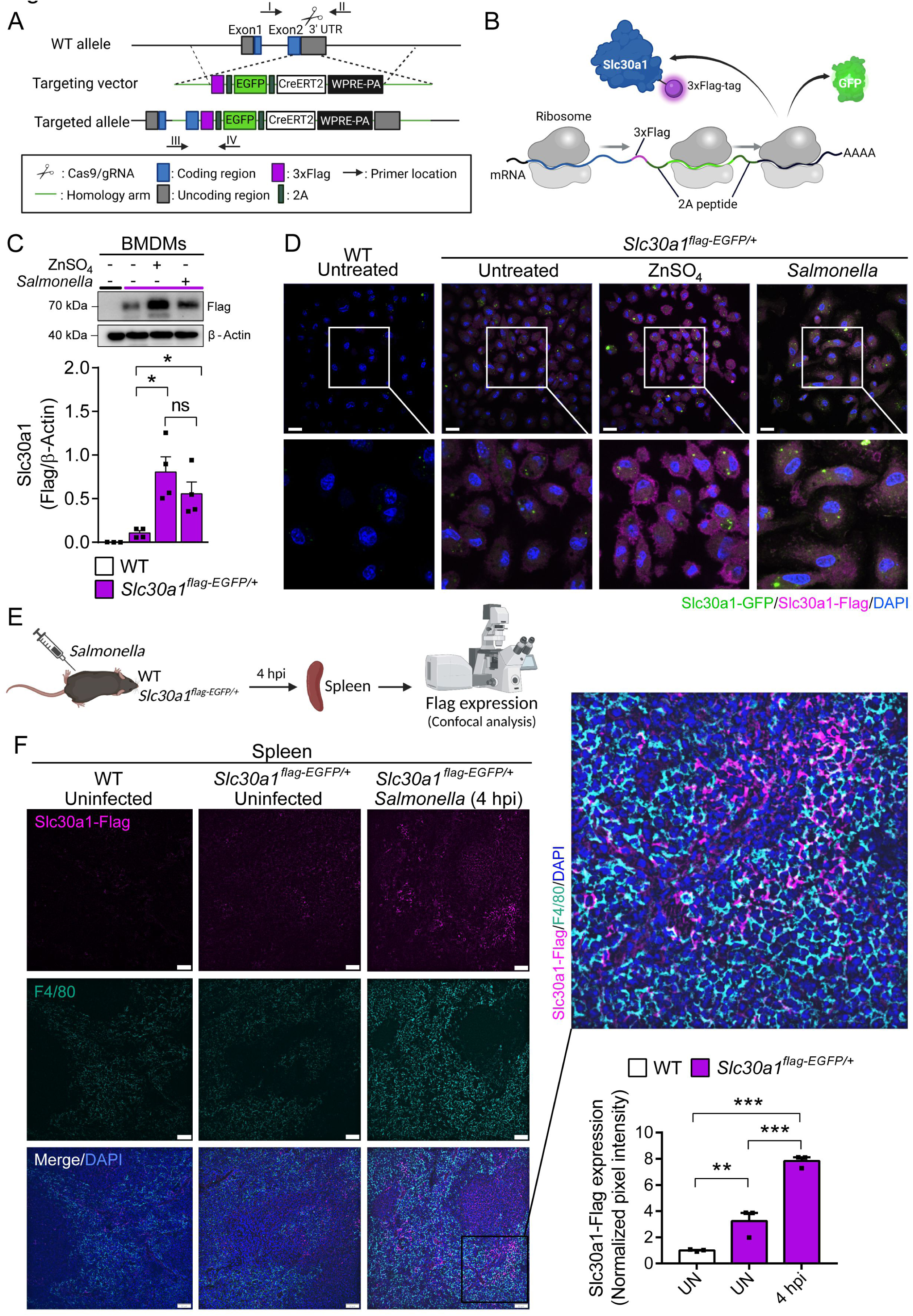
Characterization of a Slc30a1 reporter mouse with or without *Salmonella* infection. (**A**) Strategy for generating an Slc30a1 reporter mouse line expressing 3xFlag-EGFP under the control of the endogenous *Slc30a1* promotor (*Slc30a1^flag-EGFP/+^*). (**B**) Schematic diagram depicting the flag-tagged protein used to measure Slc30a1 expression and localization, with GFP serving as a transcriptional reporter cleaved by a ribosome-skipping 2A sequence on both sides of GFP and downstream of the 3xFlag sequence. (**C**) Western blot analysis of Slc30a1-flag in BMDMs isolated from *Slc30a1^flag-EGFP/+^*mice and exposed to ZnSO_4_ treatment (40 µM) or *Salmonella* (MOI = 1) for 4 h. (**D**) Confocal fluorescence images of BMDMs treated as shown in C and immunostained using an anti-flag antibody (magenta); the nuclei were counterstained with DAPI (blue), and GFP was visualized directly based on green fluorescence. Scale bars, 20 µm. (**E**) Schematic diagram depicting the strategy used to measure Slc30a1-flag expression in the spleen of *Salmonella*-infected *Slc30a1^flag-EGFP/+^* mice at 4 hpi. (**F**) Confocal fluorescence images of spleen samples obtained from a *Salmonella*-infected mouse at 4 hpi; also shown are samples of uninfected WT and *Slc30a1^flag-EGFP/+^* mice. The samples were stained with anti-flag (magenta) and anti-F4/80 (cyan), and the nuclei were counterstained with DAPI (blue). Scale bars, 50 µm. Shown at the right is the summary of the normalized pixel intensity of Slc30a1-flag expression in the spleen in each group. Data in C and F are presented as mean ± SEM. *P* values in C and F were determined using 2-tailed unpaired Student’s *t-*test. \**P*<0.05, \*\**P*<0.01 and ns, not significant.

To examine the *in vivo* expression of Slc30a1, we infected *Slc30a1^flag-EGFP/+^*mice with an i.p. injection of *Salmonella* (1×10^5^ CFU per mouse) (Figure 2E and Figure 3A). Given that the spleen is the largest secondary lymphoid organ *(Lewis et al., 2019)* and is a major source of bacterial burden during systemic infection *(Carreno et al., 2021)*, we dissected the spleen 4 h after infection (4 hpi) and examined Slc30a1-flag expression in splenic macrophages. Immunostaining for Slc30a1-flag and the macrophage biomarker F4/80 revealed that Slc30a1 is expressed at high levels specifically in splenic F4/80‒positive cells at the area of red pulp in infected mice, but not in uninfected mice (Figure 2F). Moreover, *Salmonella*-induced expression of Slc30a1 was confirmed by measuring high GFP expression in CD11b^+^F4/80^+^ peritoneal macrophages (Figure 3B) and splenic macrophages (Figure 3C, D), while there was no GFP expression in other splenic immune cells (Figure 3E-G).

**Figure 3.**
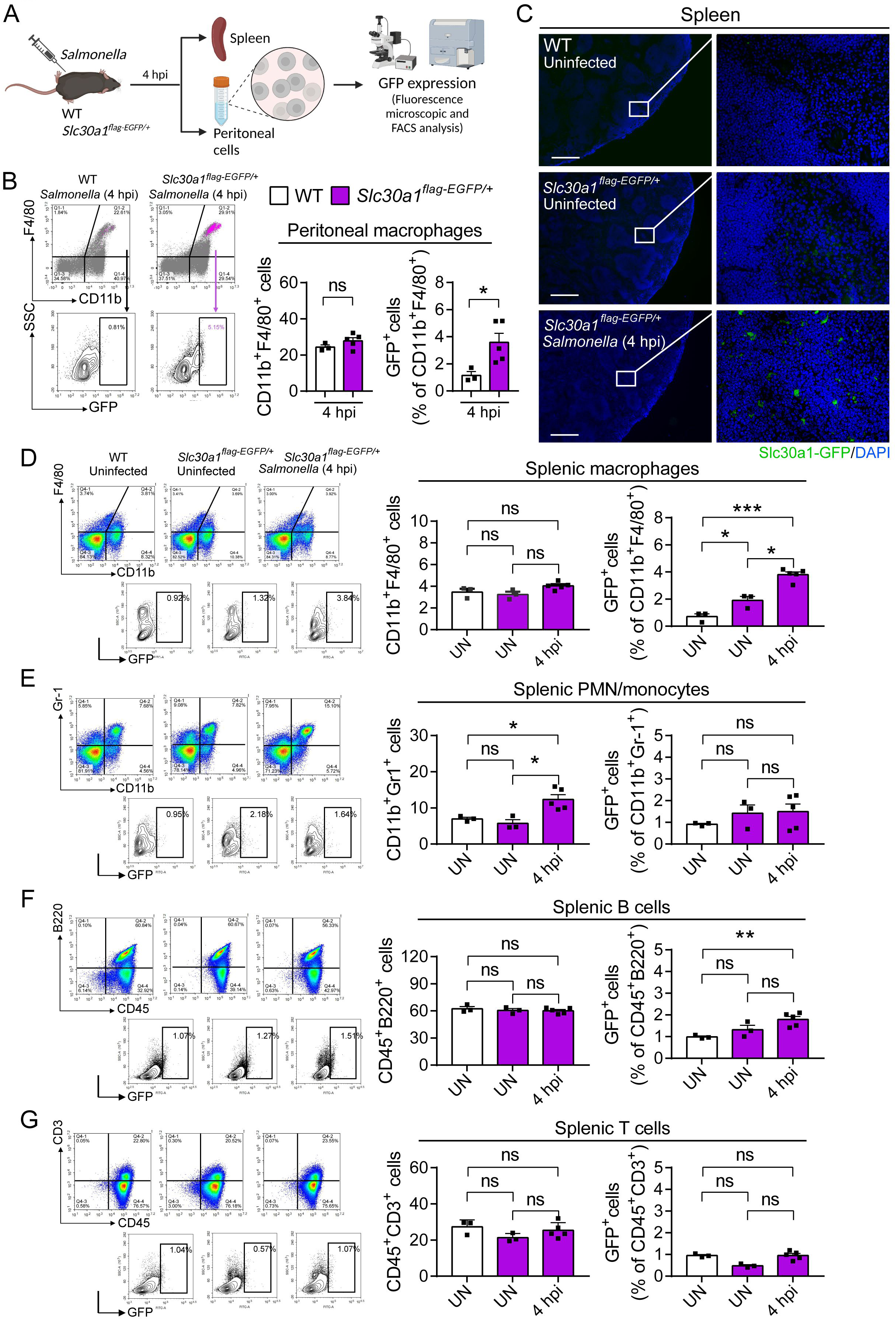
Induction of Slc30a1 expression in macrophages of a Slc30a1 reporter mouse upon *Salmonella* infection. (**A**) Schematic diagram depicting the strategy used to measure Slc30a1-GFP expression in immune cells in the peritoneal cavity and spleen of *Salmonella*-infected *Slc30a1^flag-EGFP/+^* mice at 4 hpi. (**B**) FACS plots of GFP expression in CD11b^+^F4/80^+^ macrophages isolated from the peritoneal cavity of *Salmonella*-infected WT mice and *Salmonella*-infected *Slc30a1^flag-EGFP/+^* mice at 4 hpi. Shown at the right is the summary of the percentage of CD11b^+^F4/80^+^ peritoneal macrophages in each group (*n* = 3-5 mice/group). (**C**) Fluorescence microscopy images of spleen samples obtained from a *Salmonella*-infected mouse at 4 hpi; also shown are samples of uninfected WT and *Slc30a1^flag-EGFP/+^* mice. The green region indicated the Slc30a1-GFP expression, and the blue region indicated the nuclei staining by DAPI. Scale bar, 500 μm. (**D-G**) FACS plots of GFP expression in CD11b^+^F4/80^+^ splenic macrophages (**D**), CD11b^+^Gr-1^+^ splenic PMN/monocyte (**E**), CD45^+^B220^+^ splenic B cells (**F**), and CD45^+^CD3^+^ splenic T cells (**G**) isolated from the spleens of uninfected WT mice, uninfected *Slc30a1^flag-EGFP/+^* mice, and *Salmonella*-infected *Slc30a1^flag-EGFP/+^* mice at 4 hpi (*n* = 3-5 mice/group). Shown at the right is the summary of the percentage of CD11b^+^F4/80^+^ splenic macrophages, CD11b^+^Gr-1^+^ splenic PMN/monocyte, CD45^+^B220^+^ splenic B cells, and CD45^+^CD3^+^ splenic T cells in each group. Data in this figure are represented as mean ± SEM. *P* values were determined using 2-tailed unpaired Student’s *t*-test. \**P*<0.05, \*\**P*<0.01, \*\*\**P*<0.001 and ns, not significant.

### Macrophage-specific *Slc30a1* knockout mice have increased susceptibility to *Salmonella* infection

To examine the role of Slc30a1 in macrophages with respect to protecting against *Salmonella* infection, we generated macrophage-specific *Slc30a1* knockout mice by crossing mice carrying a floxed *Slc30a1* allele (*Slc30a1^fl/fl^*) with LysM-Cre recombinase (*LysM^Cre^*) mice (Figure 4A); we then used the homozygous floxed mice with heterozygous Cre (*Slc30a1^fl/fl^LysM^Cre^*), with homozygous floxed littermates lacking Cre (*Slc30a1^fl/fl^*) serving as a control group. Loss of *Slc30a1* expression was confirmed in *Slc30a1^fl/fl^LysM^Cre^* BMDMs, with a more than 95% reduction in *Slc30a1* mRNA levels compared to *Slc30a1^fl/fl^* cells (Figure 4B). Next, we injected *Salmonella* (1×10^5^ CFU per mouse) into male *Slc30a1^fl/fl^LysM^Cre^* and *Slc30a1^fl/fl^*littermates; we then monitored survival for 2 weeks and sacrificed a subgroup of mice 4–48-hour post infection (hpi) (Figure 4C). We found that the *Salmonella*-infected *Slc30a1^fl/fl^LysM^Cre^*mice had significantly lower survival compared to *Salmonella*-infected *Slc30a1^fl/fl^* mice (0% versus 68.5% survival, respectively) (Figure 4D). At 24 hpi, we observed severe tissue damage in the spleen (i.e., deformation of white pulp tissue) and liver (i.e., necrotic tissue) of *Slc30a1^fl/fl^LysM^Cre^* mice, but not in *Slc30a1^fl/fl^*mice (Figure 4E), as well as higher levels of serum TNFα, ALT, and AST compared to *Slc30a1^fl/fl^* mice (Figure 4F-H). Interestingly, we also found that the *Salmonella*-infected *Slc30a1^fl/fl^LysM^Cre^*mice contained a smaller proportion of CD11b^+^F4/80^+^ peritoneal macrophages compared to *Salmonella*-infected *Slc30a1^fl/fl^* mice (Figure 4I). In addition, we found significantly fewer neutrophils and monocytes in blood samples obtained at 24 hpi from infected *Slc30a1^fl/fl^LysM^Cre^* mice compared to controls (Figure 4J) without interruption of other blood parameters (Table supplement 3), possibly due to the reported expression of LysM-Cre in neutrophils *(Clausen et al., 1999)*. Moreover, peritoneal macrophages are resident cells in tissues under steady-state conditions and can become peripheral blood monocytes during the inflammatory response and infection, playing an important role in bacterial clearance *(Gordon and Taylor, 2005; Wynn et al., 2013)*.

**Figure 4.**
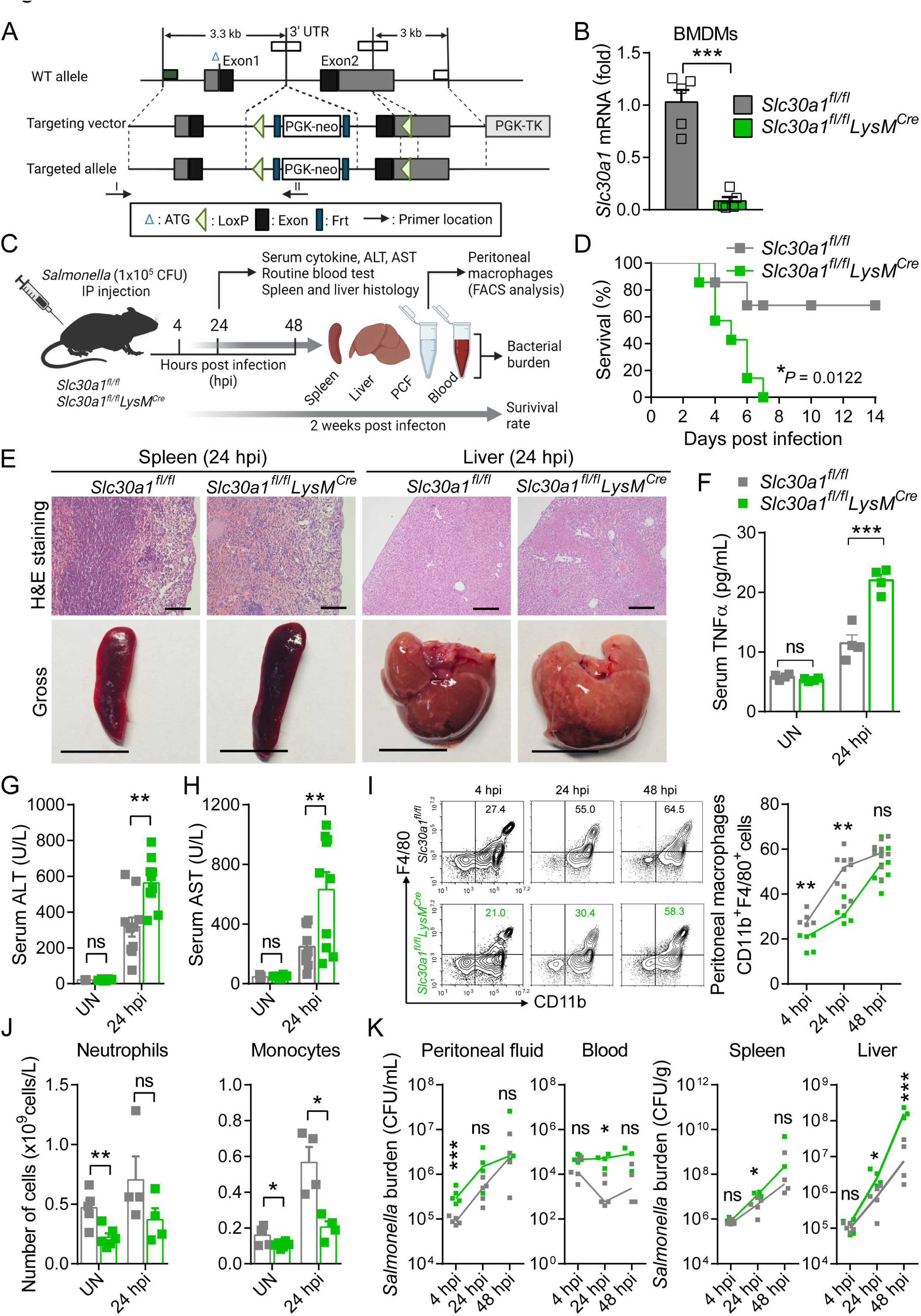
Macrophage-specific *Slc30a1* knockout mice are highly susceptible to *Salmonella* infection. (**A**) Strategy for generating macrophage-specific *Slc30a1* conditional knockout (*Slc30a1^fl/fl^*) mice. Crossing this mouse with a heterozygous *LysM^Cre^* mouse produces *Slc30a1^fl/fl^LysM^Cre^* and *Slc30a1^fl/fl^* littermates. (**B**) RT-qPCR analysis of relative *Slc30a1* mRNA levels in BMDMs isolated from *Slc30a1^fl/fl^LysM^Cre^* and *Slc30a1^fl/fl^* mice (*n* = 5). (**C**) Schematic diagram illustrating the experimental design for measuring the role of Slc30a1 in response to *Salmonella* infection. Adult (8-week-old) *Slc30a1^fl/fl^LysM^Cre^* and *Slc30a1^fl/fl^* male mice were infected by an intraperitoneal injection of 10^5^ CFU *Salmonella*. (**D**) Kaplan–Meier survival curve of *Salmonella*-infected mice (*n* = 7 mice/group). (**E**) H&E-stained and gross images of spleen and liver obtained from *Salmonella*-infected mice at 24 hpi. Scale bars, 5 mm and 10 mm, respectively. (**F-H**) Serum TNFα (**F**), ALT (**G**), and AST (**H**) were measured in uninfected (UN) and *Salmonella*-infected mice at 24 hpi (*n* = 4-10 mice/group). (**I**) FACS plots (left) and summary (right) of CD11b^+^F4/80^+^ peritoneal macrophages obtained from *Salmonella*-infected mice at 4, 24, and 48 hpi (*n* = 5-10 mice/group). (**J**) Summary of neutrophils and monocytes measured in blood samples obtained from uninfected and *Salmonella*-infected mice at 24 hpi (*n* = 4-6 mice/group). (**K**) Summary of *Salmonella* CFUs measured in the peritoneal fluid, blood, spleen, and liver of *Salmonella*-infected mice at 4, 24, and 48 hpi (*n* = 3-5 mice/group). Data in this figure are represented as mean ± SEM. *P* values of survival in D were determined using Log-rank test, in B, F, G, H, I, J and K using 2-tailed unpaired Student’s *t*-test. \**P*<0.05, \*\**P*<0.01, \*\*\**P*<0.001 and ns, not significant.

We also quantified *Salmonella* burden at 4, 24, and 48 hpi and found significantly higher numbers of bacterial cells in the peritoneal cavity, blood, spleen, and liver of infected *Slc30a1^fl/fl^LysM^Cre^*mice compared to control mice, particularly at 24 hpi (Figure 4K), indicating severe systemic infection. Furthermore, we performed *in vivo* infection studies in a separate group of mice using a non-lethal dose of *Salmonella* (1×10^4^ CFU per mouse) and found the same pattern of tissue damage (Figure supplement 2A), peritoneal macrophage recruitment (Figure supplement 2B) and bacterial burden (Figure supplement 2C). Together, these *in vivo* data provide compelling evidence that Slc30a1 in macrophages may play an important protective role against *Salmonella* infection.

### Loss of Slc30a1 in macrophages reduces bactericidal capacity by reducing iNOS and NO production

Killing intracellular microbes is a key biological function of macrophages *(Flannagan et al., 2009)*. To study whether Slc30a1 mediates this function in macrophages, we isolated BMDMs from *Slc30a1^fl/fl^LysM^Cre^*and *Slc30a1^fl/fl^* mice and exposed the cells to *Salmonella* (MOI = 10) *in vitro* for 24 h. We found that *Slc30a1^fl/fl^LysM^Cre^*BMDMs contained a significantly higher intracellular bacterial load compared to control cells at both 8 and 24 h (Figure 5A). Moreover, examining the cells at high magnification using transmission electron microscopy revealed increased numbers of bacterial cells in the phagosomes of *Slc30a1^fl/fl^LysM^Cre^*BMDMs at 24 h compared to control cells (Figure 5B).

**Figure 5.**
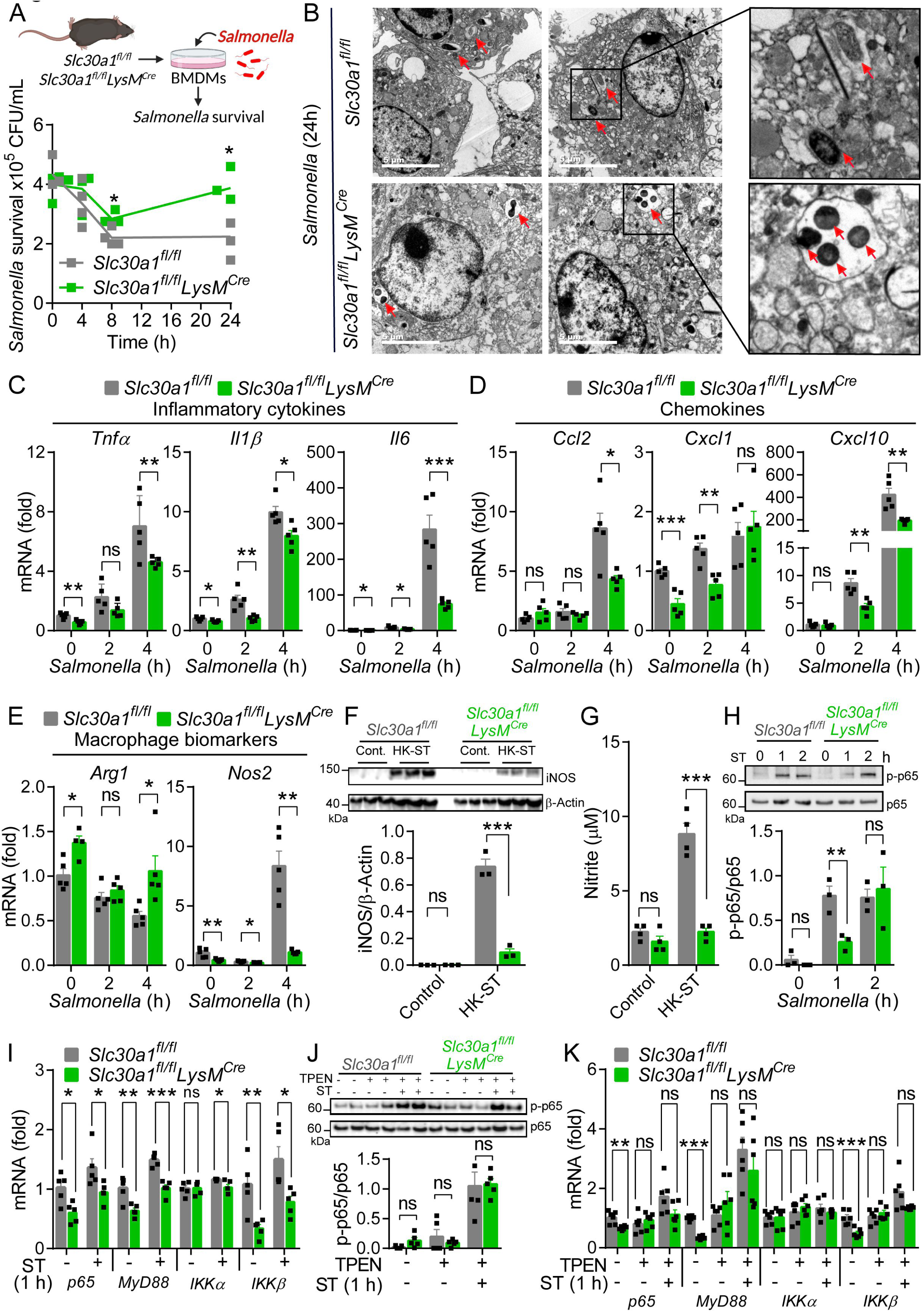
Slc30a1 is required for iNOS/NO production and intracellular pathogen-killing capacity of macrophages in response to *Salmonella* infection. (**A**) Schematic diagram (top) and time course (bottom) of the intracellular pathogen-killing capacity of *Salmonella*-infected *Slc30a1^fl/fl^LysM^Cre^* and *Slc30a1^fl/fl^* BMDMs measured in colony-forming units per ml (*n* = 3). (**B**) Transmission electron microscopy images of *Salmonella*-infected BMDMs at 24 hpi. Red arrows indicate bacterial-containing phagosomes, and the insets show magnified images of bacterial engulfment. Scale bars, 5 µm. (**C-E**) RT-qPCR analysis of mRNAs encoding inflammatory cytokines (*Tnfα*, *Il1β*, and *Il6*), chemokines (*Ccr2*, *Ccl2*, *Cxcl1*, and *Cxcl10*) and macrophage biomarkers (*Nos2* and *Arg1*) in BMDMs measured at the indicated times after *Salmonella* infection (MOI = 1) (*n* = 5). (**F**) Western blot analysis and summary of iNOS protein in *Slc30a1^fl/fl^LysM^Cre^* and *Slc30a1^fl/fl^* BMDMs either untreated or stimulated with HK-ST for 24 h (*n* = 3). (**G**) Summary of nitrite concentration measured in the cell culture supernatant of BMDMs either untreated or stimulated with HK-ST (MOI = 10) for 24 h (*n* = 4). (**H**) Western blot analysis and summary of p-p65 and p65 measured at the indicated times in *Salmonella*-infected BMDMs (*n* = 3). (**I**) RT-qPCR analysis of mRNA levels of genes involved in NF-κB signaling (*p65*, *MyD88*, *Ikkα*, and *Ikkβ*) measured in BMDMs 60 min after *Salmonella* infection (*n* = 5). (**J**) Western blot analysis and summary of p-p65 and p65 measured in uninfected and *Salmonella*-infected BMDMs either with or without TPEN (4 µM) for 60 min (*n* = 4). (**K**) RT-qPCR analysis of *p65*, *MyD88*, *Ikkα*, and *Ikkβ* mRNA in uninfected and *Salmonella*-infected BMDMs either with or without TPEN for 60 min (*n* = 6). Data in this figure are represented as mean ± SEM. *P* values in A, C, D, E, F, G, H, I, J, and K were determined using 2-tailed unpaired Student’s *t-*test. \**P*<0.05, \*\**P*<0.01, \*\*\**P*<0.001 and ns, not significant.

Infected macrophages typically exhibit a classically activated (i.e., M1-like) phenotype. During *Salmonella* infection, these activated macrophages produce several proinflammatory cytokines and chemokines such as IL-1β, TNF, and CCL2 in order to clear the pathogen via mechanisms that include cell death, the recruitment of polymorphonuclear phagocytes, and oxidative processes *(Brennan and Cookson, 2000; Pham et al., 2020; Depaolo et al., 2005)*. We therefore infected BMDMs with *Salmonella* (MOI = 1) and measured the mRNA levels of genes expressing key proinflammatory cytokines (*Tnfα*, *Il1β*, and *Il6*; Figure 5C), chemokines (*Ccl2*, *Cxcl1*, and *Cxcl10*; Figure 5D), and macrophage polarization‒related factors (*Arg1* and *Nos2*; Figure 5E). We found significantly reduced expression of proinflammatory cytokine and chemokine-related genes in infected *Slc30a1^fl/fl^LysM^Cre^* BMDMs compared to control cells. In particular, we found extremely low levels of *Nos2* mRNA in infected *Slc30a1^fl/fl^LysM^Cre^* BMDMs; *Nos2* encodes iNOS, a hallmark of classically activated macrophages that contributes to the host defense system by producing NO. In addition, we found reduced expression of *Nos2* and a number of proinflammatory cytokine-related genes in *Slc30a1^fl/fl^LysM^Cre^*BMDMs exposed to either HK-ST or LPS compared to control cells (Figure supplement 3A, B). Western blot analysis of iNOS protein levels confirmed that both HK-ST (Figure 5F) and LPS (Figure supplement 3C) downregulate *Nos2* in *Slc30a1^fl/fl^LysM^Cre^*BMDMs compared to control cells. Moreover, we measured significantly lower levels of nitrite (NO_2_^-^), a product of NO metabolism, in the cell culture supernatant of *Slc30a1^fl/fl^LysM^Cre^* BMDMs 24 h after HK-ST (Figure 5G) and LPS (Figure supplement 3D) compared to control cells.

A variety of cellular signaling pathways have been reported to regulate *Nos2* expression in macrophages in response to pathogens, particularly the NK-κB (nuclear factor kappa B) pathway *(Xie et al., 1994)*, the MAPK (mitogen-activated protein kinase) pathway *(Chan and Riches, 2001)*, and the transcription factor STAT3 (signal transducer and activator of transcription 3) *(Ahuja et al., 2020)*. We therefore measured the phosphorylated (i.e., activated) state of the principal proteins in these pathways—namely, phosphorylated p65 (in the NK-κB pathway), Erk and p38 (in the MAPK pathway), and Stat3—in *Slc30a1^fl/fl^LysM^Cre^*and *Slc30a1^fl/fl^* BMDMs 30 and 60 min after LPS stimulation. Western blot analysis revealed that p-p65, p-Erk, p-p38, and p-Stat3 levels were raised within 30 min in both *Slc30a1^fl/fl^LysM^Cre^* and *Slc30a1^fl/fl^*BMDMs; however, p-p65 levels were significantly lower in *Slc30a1^fl/fl^LysM^Cre^*BMDMs at 60 min (Figure supplement 3E). Similarly, *Salmonella* infection (MOI = 1) caused lower levels of p-p65 protein in *Slc30a1^fl/fl^LysM^Cre^* BMDMs at 60 min compared to infected *Slc30a1^fl/fl^* cells (Figure 5H), as well as reduced mRNA levels of several genes upstream of the NF-κB pathway, including *p65*, *MyD88*, *Ikkα*, and *Ikkβ* (Figure 5I). Notably, these *Salmonella*-induced changes in the expression of these key NK-κB signaling molecules was prevented by treating cells with the membrane-permeable zinc chelator TPEN (*N*,*N*,*N’*,*N’*-tetrakis (2-pyridylmethyl) ethylenediamine (TPEN)) (Figure 5J, K). These results suggest that Slc30a1 in macrophages protects against *Salmonella* infection by regulating intracellular zinc.

### Altered zinc distribution in *Salmonella*-infected macrophage-specific *Slc30a1* knockout mice

Next, we examined whether Slc30a1 in macrophages regulates systemic zinc homeostasis by measuring zinc concentration in the serum, spleen, and liver of uninfected and *Salmonella*-infected (1×10^5^ CFU per mouse) *Slc30a1^fl/fl^LysM^Cre^* and *Slc30a1^fl/fl^* mice after 24 h using inductively coupled plasma mass spectrometry (ICP-MS) (Figure 6A); we also measured other minerals, including iron, copper, magnesium, calcium, and manganese (Figure supplement 4A-C). We found that compared to *Salmonella*-infected control mice, *Salmonella*-infected *Slc30a1^fl/fl^LysM^Cre^* mice had significantly higher levels of zinc in the serum and spleen, but no difference in the liver (Figure 6B), consistent with the previous finding that during systemic infection serum zinc is redistributed to the liver in order to increase monocyte maturation, boost the immune system, and restrict the delivery of nutrient metals to pathogens *(Alker and Haase, 2018)*. In our mouse model, elevated serum zinc may impair the function of immune cells, leading to increased susceptibility to *Salmonella* infection. Moreover, the spleen serves to filter the blood filter and is a hub for monocytes *(Swirski et al., 2009)*; thus, the high zinc content in the spleen may be due to circulating high zinc‒containing macrophages, a notion supported by our finding of increased mRNA levels of *Mt1*, which encodes the principal splenic zinc reservoir protein metallothionein 1, in the spleen of *Salmonella*-infected *Slc30a1^fl/fl^LysM^Cre^*mice, but not in the liver (Figure 6C).

**Figure 6.**
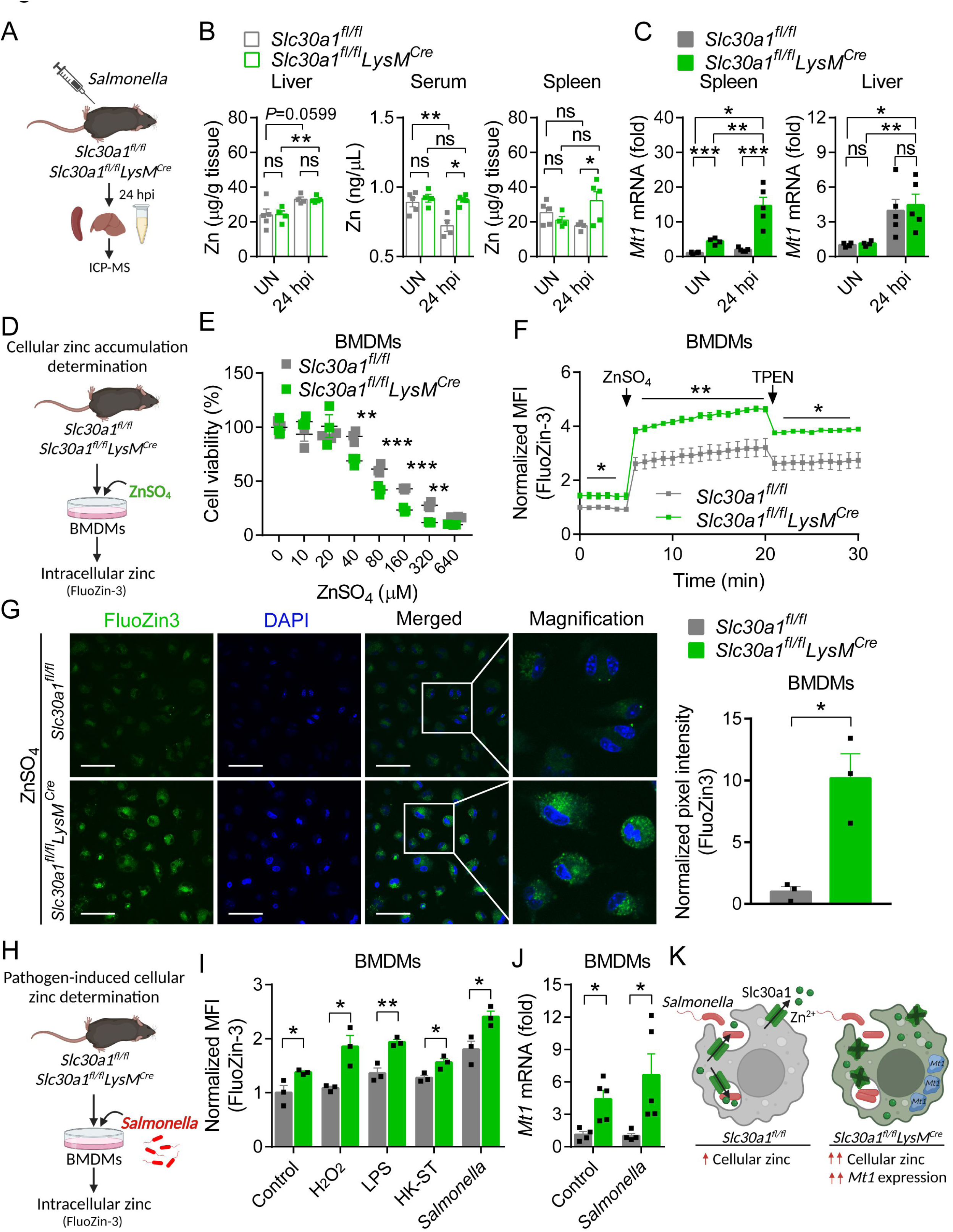
Loss of *Slc30a1* in macrophages causes intracellular zinc accumulation. (**A**) Schematic diagram depicting the strategy used to measure various trace minerals in uninfected and *Salmonella*-infected *Slc30a1^fl/fl^* and *Slc30a1^fl/fl^LysM^Cre^* mice using inductively coupled plasma mass spectrometry (ICP-MS). (**B**) Summary of zinc (Zn) content measured in the serum, spleen, and liver of uninfected (UN) and *Salmonella*-infected mice at 24 hpi (*n* = 4-5 mice/group). (**C**) RT-qPCR analysis of *Mt1* mRNA in the spleen and liver of the indicated mice. (**D**) Schematic diagram depicting the strategy used to measure intracellular zinc in BMDMs. (**E**) Time course of cell viability of *Slc30a1^fl/fl^* and *Slc30a1^fl/fl^LysM^Cre^* BMDMs after exposure to ZnSO_4_ at the indicated concentrations for 24 h. (**F**) Time course of normalized FluoZin-3 mean fluorescence intensity (MFI) measured in BMDMs; where indicated, ZnSO_4_ (100 µM) and TPEN (4 µM) were applied to the cells. (**G**) Confocal fluorescence images of BMDMs after treatment with ZnSO_4_ for 15 min; the nuclei were counterstained with DAPI (blue). Scale bars, 50 µm. Shown at the right is the summary of the normalized pixel intensity of FluoZin-3 in BMDMs. (**H**) Same as D, except the cells were infected with *Salmonella*. (**I**) Summary of normalized FluoZin-3 MFI measured in BMDMs 30 min after application of H_2_O_2_ (1 mM), LPS (1 µg/ml), HK-ST (MOI = 100), or *Salmonella* (MOI = 10) (*n* = 3). (**J**) RT-qPCR analysis of *Mt1* mRNA in uninfected and *Salmonella*-infected BMDMs (*n* = 5). (**K**) Model showing the predicted effects of the loss of Slc30a1 on cellular zinc trafficking and intracellular zinc accumulation in BMDMs in response to *Salmonella* infection. Data in this figure are represented as mean ± SEM. *P* values in B, C, E, F, G, I, and J were determined using 2-tailed unpaired Student’s *t-*test. \**P*<0.05, \*\**P*<0.01, \*\*\**P*<0.001 and ns, not significant.

Next, we used the fluorescent zinc probe FluoZin-3 to quantify the accumulation of intracellular zinc in BMDMs isolated from *Slc30a1^fl/fl^LysM^Cre^*and control mice following ZnSO_4_ treatment (Figure 6D). Consistent with the well-documented role of Slc30a1 in zinc resistance *(Palmiter, 2004)*, we found that *Slc30a1^fl/fl^LysM^Cre^*BMDMs were significantly more sensitive to ZnSO_4_-induced toxicity compared to control cells (Figure 6E), and ZnSO_4_ treatment (100 µM) caused a significantly higher FluoZin-3 signal in *Slc30a1^fl/fl^LysM^Cre^*BMDMs compared to control cells (Figure 6F, G).

Previous studies showed that cytosolic free zinc increases in macrophages following exposure to LPS, *E. coli*, and *Salmonella (Haase et al., 2008; Brieger et al., 2013; Wu et al., 2017)*. We therefore measured intracellular free zinc in *Salmonella*-infected *Slc30a1^fl/fl^LysM^Cre^*and control BMDMs (Figure 6H). We found that 30 min after *Salmonella* infection (MOI = 1), FluoZin-3 fluorescence was significantly higher in *Slc30a1^fl/fl^LysM^Cre^*cells; similar results were obtained when the cells were exposed to H_2_O_2_ (1 mM), LPS (1 µg/ml), or HK-ST (MOI = 100) (Figure 6I). In addition, we measured higher levels of *Mt1* mRNA in *Slc30a1^fl/fl^LysM^Cre^* BMDMs compared to *Slc30a1^fl/fl^*cells, even in uninfected cells (Figure 6J). Based on these results, we hypothesize that the loss of Slc30a1 leads to an abnormal increase in intracellular zinc during *Salmonella* infection and increases the expression of *Mt1*, possibly to compensate for the excessive intracellular zinc by increasing Mt1-mediated zinc storage (Figure 6K).

### Slc30a1 and Mt1 in macrophages coordinate *in vivo* survival in response to *Salmonella* infection

We hypothesized that the overexpression of Mt1 can provide an excess source of cellular zinc from which free zinc can be released via the redox reaction during pathogen-induced oxidative stress. We therefore asked whether knocking out *Mt1* in Slc30a1-deficient macrophages can reduce intracellular zinc and restore their antimicrobial response. To test this notion, we generated double-knockout (*Slc30a1^fl/fl^LysM^Cre^;Mt1^-/-^*) mice (referred to hereafter as DKO mice) by crossing *Slc30a1^fl/fl^LysM^Cre^* mice with *Mt1^-/-^*mice and then isolated BMDMs, with *Slc30a1^fl/fl^* and *Slc30a1^fl/fl^;Mt1^-/-^*BMDMs serving as controls (Figure 7A, B). We first determined whether NF-κB signaling in BMDMs could be restored by the loss of Mt1. Both western blot and RT-qPCR analyses revealed that p-p65 protein levels (Figure 7C) and *p65*, *MyD88*, *Ikkα*, and *Ikkβ* mRNA levels (Figure 7D) were similar to control cells in *Salmonella*-infected DKO cells. Strikingly, however, DKO BMDMs had the lowest killing capacity among all four groups (Figure 7E). In addition, the concentration of nitrite in the cell culture supernatant was significantly lower in DKO BMDMs 24 h after LPS and HK-ST stimulation (Figure 7F). We then measured intracellular zinc levels using FluoZin-3 and found similar levels between DKO and *Slc30a1^fl/fl^LysM^Cre^* BMDMs 30 min after stimulation with either LPS or HK-ST (Figure 7G). Finally, following a 4-hour pretreatment with ZnSO_4_ (40 µM), we measured higher levels of zinc in DKO BMDMs compared to *Slc30a1^fl/fl^LysM^Cre^* BMDMs (Figure 7H), suggesting that Mt1 may indeed play a role in maintaining the labile zinc pool.

**Figure 7.**
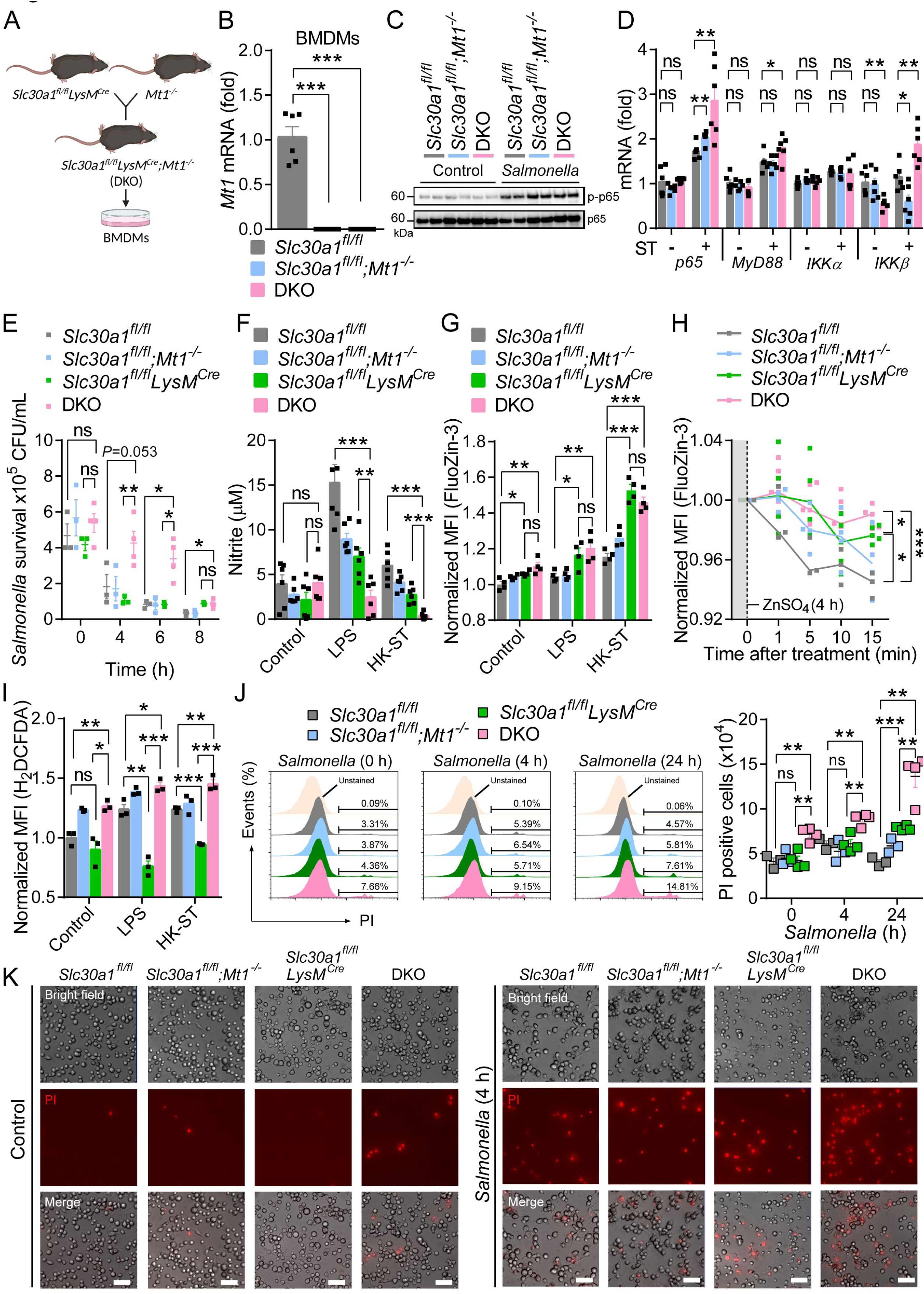
Slc30a1 and Mt1 coordinate macrophage survival in response to *Salmonella* infection. (**A**) Breeding strategy for generating *Slc30a1* and *Mt1* double-knockout (*Slc30a1^fl/fl^LysM^Cre^; Mt1^-/-^*) BMDMs (DKO). (**B**) Summary of *Mt1* mRNA measured in *Slc30a1^fl/fl^*, *Slc30a1^fl/fl^; Mt1^-/-^*, and DKO BMDMs (*n* = 6). (**C**) Western blot analysis and summary of p-p65 and p65 measured in the indicated BMDMs 60 min after *Salmonella* infection (MOI = 1). (**D**) RT-qPCR analysis of *p65*, *MyD88*, *IKKα*, and *IKKβ* mRNA measured in uninfected and *Salmonella*-infected BMDMs (*n* = 6). (**E**) Time course of *Salmonella* CFUs measured in the indicated BMDMs (*n* = 3-4). (**F**) Summary of nitrite concentration in the cell culture supernatant of untreated BMDMs and BMDMs 24 h after LPS (100 ng/ml) or HK-ST (MOI = 10) stimulation (*n* = 6). (**G**) Summary of intracellular zinc measured in untreated BMDMs and BMDMs 30 min after LPS (1 µg/ml) or HK-ST (MOI = 100) stimulation (*n* = 4). (**H**) Time course of intracellular zinc measured in BMDMs after a 4-h pretreatment (shaded area) with ZnSO_4_ (40 µM). (**I**) Summary of intracellular ROS concentration measured using H_2_DCFDA staining in untreated BMDMs and 30 min after stimulation with LPS (1 µg/ml) or HK-ST (MOI = 100) (*n* = 3). (**J**) FACS plots (left) and summary (right) of propidium iodide (PI)‒positive BMDMs at the indicated times after *Salmonella* infection (MOI = 10) (*n* = 3-4). (**K**) Fluorescence microscopy images of PI-stained BMDMs either uninfected (left) or 4 h after *Salmonella* infection (right). Scale bars, 50 µm. Data in this figure are represented as mean ± SEM. *P* values in B, D, E, F, G, H, I, and J were determined using 2-tailed unpaired Student’s *t-*test. \**P*<0.05, \*\**P*<0.01, \*\*\**P*<0.001 and and ns, not significant.

The metal-binding protein Mt1 is responsible for metal detoxification and storage, as well as helping prevent cellular stress from oxidation and inflammation *(Dai et al., 2021)*. Indeed, excess zinc can act as a pro-oxidant *(Lee, 2018)* that triggers cell death *(Franklin and Costello, 2009)*. To address the possibility that the excess zinc caused by the loss of Mt1 can increase cellular stress and induce cell death, we used the fluorescent probe H_2_DCFDA to measure the production of reactive oxygen species (ROS) and then performed propidium iodide (PI) staining to measure cell death. As expected, DKO BMDMs stimulated with either LPS or HK-ST had increased ROS production at 30 min compared to *Slc30a1^fl/fl^* cells, whereas ROS levels were significantly reduced in *Slc30a1^fl/fl^LysM^Cre^*BMDMs compared to *Slc30a1^fl/fl^* cells (Figure 7I). Consistent with this increase in ROS production in DKO BMDMs, we found a significant increase in PI-positive DKO BMDMs both 4 and 24 h after *Salmonella* infection (Figure 7J, K) as well as LPS treatment (Figure supplement 5).

## Discussion

Here, we provide evidence that *Slc30a1* expression in macrophages is required for host resistance against pathogens, illustrated schematically in Figure 8. Based on this model, *Slc30a1*-deficient macrophages have increased *Salmonella* burden due to reduced iNOS and NO formation via reduced NF-κB signaling. This effect is due primarily to the inhibitory effects of high intracellular zinc, reflected by the compensatory upregulation of the zinc-binding protein Mt1. However, knocking out *Mt1* in *Slc30a1*-deficient macrophages does not restore bactericidal activity; rather, the loss of Mt1 results in extremely high zinc accumulation under pathogen stimulation, engaging oxidative stress pathways and leading to increased cell death.

**Figure 8.**
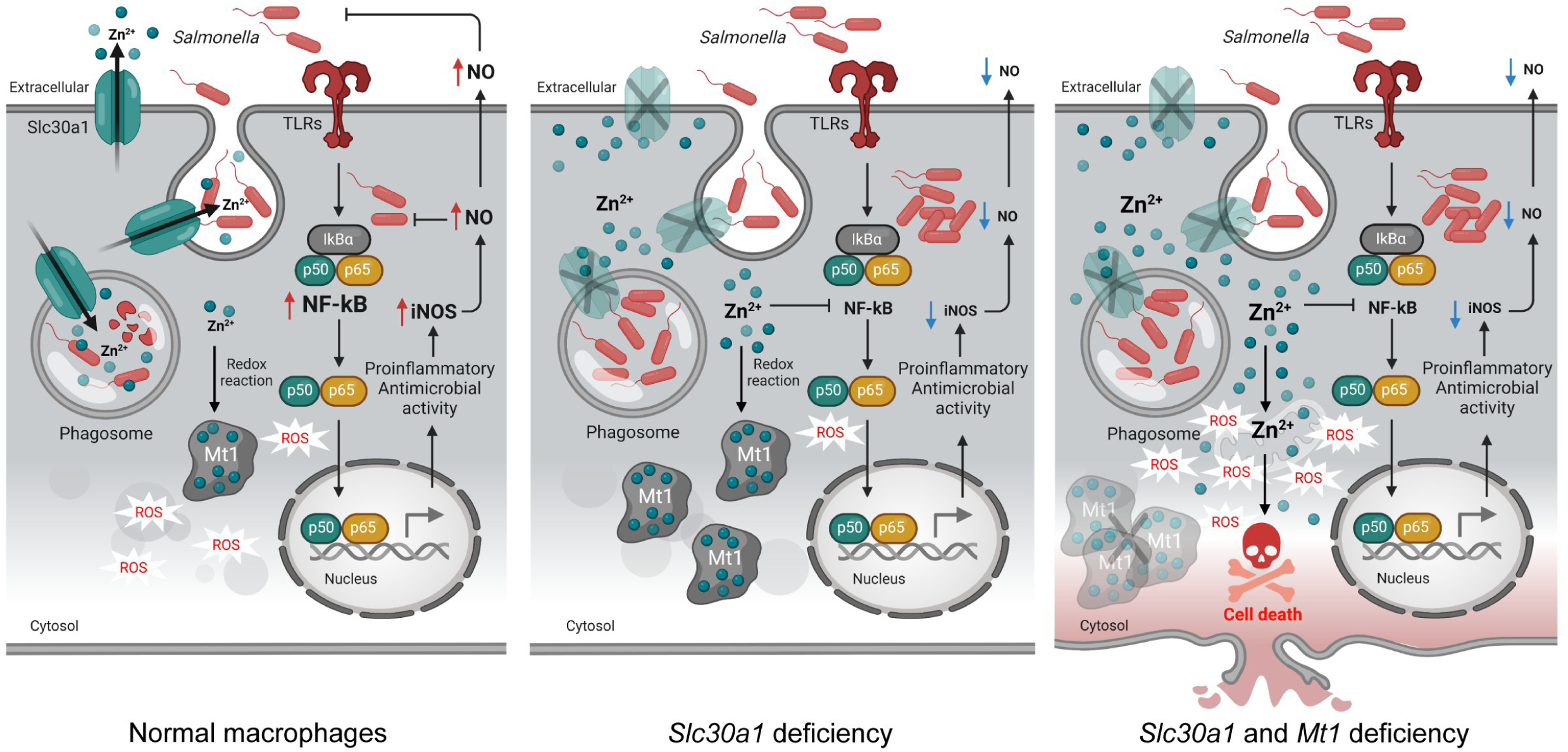
Putative protective function of Slc30a1 in macrophages during *Salmonella* infection. Left, under normal conditions, *Slc30a1* is upregulated in response to *Salmonella* infection, generating a short-term decrease in cytosolic zinc concentration and increasing zinc toxicity in *Salmonella*-containing phagosomes. Middle, loss of Slc30a1 leads to an accumulation of intracellular zinc, thereby upregulating *Mt1* overexpression and reducing iNOS and NO production via reduced NF-κB signaling, reducing the cell’s bacterial clearance capacity. Right, Mt1 is a key zinc-binding protein that helps buffer excess zinc. Loss of both Mt1 and Slc30a1 leads to increased ROS formation, resulting in cell death due to an inability to protect against intracellular zinc saturation.

The cellular zinc exporter Slc30a1 is a key regulator of cellular zinc homeostasis *(Kambe et al., 2015; Liuzzi and Cousins, 2004; Lichten and Cousins, 2009)*. Previous *in vitro* studies have shown that Slc30a1 in macrophages mediates the intracellular killing of invading *Mycobacterium tuberculosis* and *E. coli* by increasing zinc levels within phagosomes, thereby protecting the cell against infection *(Botella et al., 2011; Stocks et al., 2021)*. Here, we provide the first *in vivo* evidence that Slc30a1 in macrophages plays a key role in host protection against *Salmonella* infection. Specifically, we show that mice lacking Slc30a1 in macrophages are highly susceptible to *Salmonella* infection. Moreover, loss of Slc30a1 in macrophages reduced the number of circulating neutrophils and monocytes and the number of peritoneal macrophages. Although the connection between Slc30a1-deficient macrophages and the proportion of these phagocytes remains unknown, it may play a role in reducing bacterial clearance, thereby contributing to the host’s increased susceptibility to *Salmonella* infection.

The killing capacity of macrophages has been studied extensively and mediates the host defense by ingesting and then destroying invading microbes *(Flannagan et al., 2009)*. Using BMDMs isolated from our macrophage-specific *Slc30a1* knockout mice, we show that these macrophages have reduced pathogen-killing capacity via reduced NF-κB signaling decreasing iNOS and NO production. Interestingly, we found that treating these cells with the zinc chelator TPEN improves NF-κB signaling, suggesting that changes in cellular zinc levels likely contribute to their reduced antimicrobial function. Wu *et al*. previously reported that zinc can suppress iNOS/NO production by inhibiting NF-κB in *Salmonella*-infected macrophages *(Wu et al., 2017)*. Moreover, recent studies have shown that zinc can reduce NF-κB activation in macrophages *(Haase et al., 2008; Brieger et al., 2013; Wu et al., 2017; Jeon et al., 2000)*. With respect to the underlying mechanism, studies have shown that zinc inhibits the NF-κB pathway by reducing the activity of recombinant IKKα and IKKβ by blocking IκBα phosphorylation and degradation *(Jeon et al., 2000; Zhou et al., 2004; Liu et al., 2013)*. Consistent with this notion, our finding of the role for Slc30a1 in regulating the macrophage response to *Salmonella* may help explain how the genetic deletion of *Slc30a1* specifically in this professional phagocyte leads to the observed disease outcome in our mouse model.

An interesting finding from our *in vivo* experiments is that our macrophage-specific *Slc30a1* knockout mice fail to redistribute zinc levels in the serum and spleen following *Salmonella* infection. During systemic infection, serum zinc is rapidly taken up by the liver as part of the host defense process *(Alker and Haase, 2018)*. In mice, defects in this process—for example, due to high zinc supplementation—have negative consequences for neutrophil extracellular traps (NETs) *(Kuźmicka et al., 2020)*. In infants, consuming a full-fat diet of powdered cow’s milk containing excessive zinc content can lead to a decreased phagocytic capacity of monocytes *(Schlesinger et al., 1993)*. In adults, excessive consumption of zinc supplements can impair both the migration and bactericidal capacity of polymorphonuclear leukocytes *(Chandra, 1984)*. This is consistent with our finding that maintaining serum zinc levels in our macrophage-specific *Slc30a1* knockout mice affects the percentage of neutrophils and monocytes and can interfere with the migration of macrophages, thereby reducing bacterial clearance and increasing their susceptibility to *Salmonella* infection. Furthermore, nutrient-dense conditions such as excessive serum zinc can be beneficial to the growth of *Salmonella*, allowing the pathogens to overcome the host’s defenses.

Our measurements of intracellular zinc levels revealed high zinc concentrations in Slc30a1-deficient BMDMs. We also measured an upregulation of *Mt1* in these cells based on our *in vivo* experiments, possibly indicating a cellular response to intracellular zinc saturation. Mt1 is a low molecular weight cysteine-rich protein essential for buffering surplus zinc in order to maintain zinc homeostasis *(Krężel and Maret, 2017)*. Interestingly, Malavolta *et al*. previously reported that transgenic mice overexpressing *Mt1* had a significantly prolonged life span compared to control mice when fed a high-zinc diet *(Malavolta et al., 2012)*. In addition, Smith *et al*. showed that an epithelial cell line overexpressing *Mt1* was less susceptible to zinc toxicity without affecting cell cycle progression, and the level of *Mt1* expression was positively correlated with intracellular free zinc concentration *(Smith et al., 2008)*. In this respect, *Mt1* expression may serve as an interesting target for reducing cellular zinc. Surprisingly, we found that macrophages lacking both Mt1 and Slc30a1 have severely reduced bactericidal capacity; moreover, these double-knockout macrophages contain higher levels of free zinc, which may partially explain their decreased antimicrobial activity.

Zinc is an essential trace element that helps reduce cellular inflammation; paradoxically, however, zinc overload can increase ROS production, leading to mitochondrial damage and reduced cellular ATP levels *(Dineley et al., 2003)*. Previous studies found that increasing intracellular zinc by inducing the lysosomal zinc channel TRPML1 causes mitochondrial dysfunction and ATP depletion, leading to necrotic cell death *(Du et al., 2021)*. In addition, metallothionein has been reported to play a protective role against zinc toxicity *(Palmiter, 2004)*. Here, we show that in the absence of Slc30a1-mediated zinc efflux and Mt1-mediated redox buffering, cytosolic free zinc increases significantly, leading to ROS overproduction and therefore presumably causing macrophage death. Moreover, in the context of pathogenesis, invading *Salmonella* may utilize excess cellular zinc as a nutrient source in order to evade the host’s antimicrobial defense, while intracellular *Salmonella* can secrete virulence proteins that increase host cell death *(Hersh et al., 1999; Hernandez et al., 2003)*.

Although we found that genetically deleting *Slc30a1* in macrophages reduces their antimicrobial capacity via the NF-κB pathway due to increased intracellular zinc, we cannot rule out the possibility that the increased intracellular zinc affects other zinc-binding proteins such as the zinc-finger protein A20 *(Prasad et al., 2011)*, which also reduces NF-κB signaling *(Boone et al., 2004; Shembade et al., 2010; Wertz et al., 2004)*. In addition, our results obtained using *Slc30a1* and *Mt1* double-knockout macrophages are consistent with increased zinc-mediated oxidative stress leading to increased ROS production and cell death. Nevertheless, the underlying mechanism that connects increased intracellular zinc, increased ROS production, and cell death remains unknown. Finally, although we focused our study on BMDMs, macrophages may response differently to infection in the context of the full immune system *in vivo*. Thus, generating a mouse line in which both *Mt1* and *Slc30a1* are knocked out specifically in macrophages may provide insight into the mechanism by which zinc-mediated macrophage cell death in the absence of both Slc30a1 and Mt1 affects disease outcome.

In conclusion, we report that Slc30a1 in macrophages plays a protective role against *Salmonella* infection by regulating the antimicrobial response and promoting cell survival by maintaining cellular zinc homeostasis. These findings underscore the notion that cellular zinc is tightly regulated in order to main immune function, thus maximizing antimicrobial activity while minimizing cellular oxidative damage. These results provide new insights into how high levels of intracellular zinc can increase the risk of impaired immune cell function, thereby affecting the host’s front line of defense against invading pathogens. From a clinical perspective, balancing zinc intake is an important step in limiting morbidity due to *Salmonella* infection.

## Materials and methods

### Mouse strains

*Slc30a1^flag-EGFP/+^* mice were generated by Shanghai Biomodel Organism Science & Technology Development Co. Ltd. The donor vector contains the 5’ homologous arm, 3xFlag-2A-EGFP-2A-CreERT2-Wpre-pA, and the 3’ homologous arm. In the 3’ UTR of the *Slc30a1* locus, the 3xFlag-2A-EGFP-2A-CreERT2-Wpre-pA cassette was inserted downstream of exon 2. *Slc30a1* floxed mice were generated by Shanghai Biomodel Organism Science & Technology Development Co. Ltd. The floxed *Slc30a1* allele contains a single loxP site upstream of exon 2 and a single loxP site neo cassette downstream of exon 2. Heterozygous *Slc30a1* floxed mice (*Slc30a1^fl/+^*) were backcrossed to the C57BL/6 background for more than 5 generations, and then crossed with LysM-Cre expressing mice (The Jackson Laboratory, 004781) to generate macrophage-specific *Slc30a1* knockout (*Slc30a1^fl/fl^LysM^Cre^*) mice. All mice were genotyped using genomic PCR. Cre-negative floxed mice (*Slc30a1^fl/fl^*) were used as the control group. Under standard conditions, 8-weeks-old male *Slc30a1^fl/fl^LysM^Cre^*present with no obvious phenotype (data not shown). *Mt1^-/-^* mice were purchased from Shanghai Biomodel Organism Science & Technology Development Co. Ltd. and crossed with *Slc30a1^fl/fl^LysM^Cre^*mice to generate double-knockout (*Slc30a1^fl/fl^LysM^Cre^; Mt1^-/-^*) mice. Wild-type C75BL/6 mice were purchased from Shanghai SLRC Laboratory Animal Co., Ltd. Eight-week-old male mice were used in this study. All animals were fed a standard chow diet with access to drinking water and were housed at constant temperature (23°C) under a 12:12-h light/dark cycle in specific pathogen-free conditions. All *in vivo* experiments were conducted in accordance with the National Institutes of Health guidelines and were approved by the Institutional Animal Care and Use Committee, Zhejiang University.

### DNA isolation and genotyping

Genomic DNA of mouse tissue biopsies was extracted using the Tissue & Blood DNA Extraction kit-250prep (Zhejiang Easy-Do Biotech CO., LTD, #DR0301250) in accordance with the manufacturer’s instructions. DNA samples were quantified using a Nanodrop 2000 spectrophotometer (Thermo Fisher Scientific). PCR amplification was conducted using 50 ng DNA per reaction, 2xTaq PCR StarMix (GenStar, #A012-101), and specific primers (Table supplement 4) in a T100 Thermal Cycler (Bio-Rad, #10878655). Finally, the PCR products were separated by agarose gel electrophoresis to determine the genotypes.

### Bacterial preparation

*Salmonella enterica* serovar Typhimurium *(Wu et al., 2017)* was grown in tryptic soy broth (TSB) medium (Solarbio Life Science, #T8880) overnight at 37°C under sterile conditions. Bacterial inoculum was compared to a 0.5 McFarland turbidity standard (approximately 1×10^8^ CFU/ml) and adjusted to 10^5^ colony forming units per ml (CFU/ml) with sterile phosphate-buffered saline (PBS). The bacterial count in the original suspension was verified using a plate counting method. Heat-killed *Salmonella typhimurium* (HK-ST) was prepared by incubating the bacterial inoculum in a 60°C water bath for 2 h, and bacterial viable count was performed to confirm ≥99.99% reduction in viability.

### Primary cell culture

Bone-marrow derived macrophages (BMDMs) were differentiated from bone marrow cells obtained from 8-week-old male mice (Figure supplement 6). In brief, bone marrow cells from femur and tibia were isolated by flushing with sterile PBS (Corning, #21-040-CV) and filtered through a 40-μm nylon membrane filter (Falcon, #350). The resulting cell suspension was centrifuged at 300x*g* for 5 min, and the cell pellet was resuspended in RPMI 1640 (Corning, #21-040-CVR) containing 1% (v/v) penicillin-streptomycin (HyClone, #SV30010), 20% (v/v) fetal bovine serum (FBS) (Gibco, #10270-106), and 30% L929 conditioned medium. BMDMs were then differentiated for 7 days under 37°C in humidified air containing 5% CO_2_ with refreshing the differentiation medium every two days. These BMDMs were then transferred to RPMI 1640 supplemented with 1% penicillin-streptomycin and 10% FBS at 37°C and used the next day. We found no difference in BDMD differentiation between the various genotypes used in this study.

### RNA-seq and data analysis

BMDMs from biological triplicates of C75BL/6 mice were infected with *Salmonella* for 2 h, and total RNA was extracted for RNA-seq. RNA-seq analysis was performed by LC Sciences (Hangzhou, China). Gene Ontology (GO) and Kyoto Encyclopedia of Genes and Genomes (KEGG) enrichment analyses were performed using the DAVID database. Heatmaps were generated using the OmicStudio tool (https://www.omicstudio.cn/tool). Volcano plots and bar plots of the GO and KEGG pathways were generated using the Tableau (2019.3. Ink) software desktop system.

### Immunofluorescence

BMDMs grown on glass coverslips in 6-well plates were exposed to ZnSO_4_ (40 µM) or infected with *Salmonella* (MOI = 1) for 4 h. The cells were then fixed in 4% paraformaldehyde (PFA) for 15 min, permeabilized in 0.2% Triton-X 100 (Sigma, #T9284) for 10 min, and then blocked in 5% (w/v) BSA (Sigma, #A6003) for 30 min at room temperature. The fixed cells were then incubated in monoclonal anti-FLAG M2-Peroxidase (HRP) antibody (Sigma, #A8592) overnight at 4°C, followed by fluorophore-conjugated secondary antibody, Alexa Fluor® 647 secondary antibody (Thermo Fisher Scientific, #A32728) for 1 h at room temperature in the dark. The nuclei were stained with 3 µM DAPI (4’,6-diamidino-2-phenylindole; BioLegend, #422801) for 5 min; after gently rinsing three times with PBS, the glass coverslips were mounted cell-side down on clean plain glass slides with 80% glycerol (Sigma, #G5516) and imaged using a CSU-W1 confocal microscope (Olympus).

### *Salmonella* infection in vivo

Eight-week-old male mice were injected intraperitoneally (i.p.) with 200 μl PBS containing *Salmonella* at a final dose of 1×10^5^ CFU per mouse or sterile PBS as uninfected groups. At the indicated time points, the mice were euthanized, peritoneal cavity cells were subsequently collected for determination of peritoneal macrophage population. Blood samples were obtained via cardiac puncture, and the spleen and liver were dissected. The organs were either immediately placed on ice in sterile PBS to count bacterial burden or immersed in 4% PFA for later histological analysis. A portion of blood was kept in serum collection tubes and allowed to clot for 2 h at room temperature; clear serum was then obtained by centrifugation at 1000 rpm for 10 min. For survival experiments, after infection with *Salmonella* the mice were monitored daily for 2 weeks.

### Bacterial burden in tissues

Approximately 100-200 mg of liver and spleen samples were homogenized in sterile PBS. A 100-µl of tissue homogenate, peritoneal cavity fluid, and blood were serially diluted 10-fold in PBS and plated on TSA agar (Solarbio life science, #T8650). The plates were incubated at 37°C overnight, and viable bacteria colonies were counted as CFU/ml or CFU per gram of tissue (CFU/g).

### Serum ALT and AST measurements

The levels of serum aspartate aminotransferase (AST) and alanine aminotransferase (ALT) from mice infected with or without *Salmonella* for 24 h were measured using alanine aminotransferase assay kit and aspartate aminotransferase assay kit (ShenSuoYouFu) according to the manufacturer’s protocol.

### ELISA

Cytokine levels of TNFα was quantified in serum obtained from mice infected with and without *Salmonella* for 24 h by the mouse/rat TNF-A Valukine ELISA kit (R&D Systems, #VAL609) following the product protocol.

### Histopathology

Spleen and liver samples were immersed in 4% PFA solution for 24 h and then embedded in paraffin. The tissue samples were then sectioned and stained with hematoxylin and eosin (H&E) using standard protocols. The sections were examined using bright-field microscopy (Nikon Eclipse Ni-U). Three adjacent sections of each sample were quantified using ImageJ software (National Institutes of Health).

### Flow cytometry

Cell suspensions (1×10^5^ cells/ml) were immunostained using specific anti-mouse antibodies in the dark at 4°C for 30 min. The following fluorochrome-conjugated antibodies were used in this study (all from BioLegend): CD11b-Pacific Blue (clone M1/70, #101224), CD11b-PE (clone M1/70, #1101208), F4/80-APC (clone BM8, #123116), Gr-1-APC (clone RB6-8C5, #108412), CD45-APC/Cyanine7 (clone 30-F11, #103116), CD45R/B220-Pacific Blue (clone RA3-6B2, #103227), and CD3-PerCP/Cyanine 5.5 (clone 17A2, #100218). All antibodies were used at a dilution of 1:200. Live versus dead cells were identified based on DAPI staining. Cell sorting were performed using the NovoCyte flow cytometer (ACEA Biosciences Inc.).

### Trace mineral analysis using ICP-MS

Metal content was measured using inductively coupled plasma mass spectrometry (ICP-MS). Tissue samples (200 mg) were digested in 4 ml of EMSURE ISO nitric acid 65% solution (Merck, #1.00456.2508) using MARS 6 microwave extraction system. In brief, the samples were heated from room temperature to 200°C in a microwave oven (1,000 W, 2,450 MHz) for up to 20 min. The microwave power was maintained for an additional 30 min followed by a cooling-down period of 15 min. The samples were then completely dried at 110°C for 3 h. The digested samples were diluted to a final volume of 5 ml in high-purity deionized water obtained using a Milli-Q Integral 10 purification system (EMD Millipore, #C205110). The resulting samples were subjected to metal analysis using an Agilent 7700x ICP-MS equipped with an Agilent ASX 520 auto-sampler.

### Gene expression analysis using RT-qPCR

Total RNA was extracted from the samples using TransZol Up (TransGen Biotech, #ET111-01), and 1 μg of total RNA was used as a template to synthesize cDNA using the HiScript II 1st Strand cDNA Synthesis Kit (Yeasen Biotech, #11123ES60) in a ProFlex PCR System (Life technologies). Each RT-qPCR reaction consisted of 5 μl of SYBR Green PCR Master Mix (Bimake, #B21202), 2 μl of forward and reverse primers (1 µM), 0.2 µl of cDNA, and sufficient RNase-free water to yield a final volume of 10 μl. RT-qPCR was performed in duplicate for each sample using the LightCycler 480 Real-Time PCR System (F. Hoffmann-La Roche, Ltd.) with the following conditions: initial denaturation at 95°C for 3 min, 40 cycles of amplification at 95°C for 15 s, 60°C for 30 s, and 72°C for 30 s, followed by denaturation at 95°C for 30 s, and annealing-extension at 40 °C for 30 s. Data were analyzed using the 2^-△△Ct^ method *(Pfaffl, 2001)* to calculate the relative target gene expression normalized to the endogenous housekeeping gene *Gapdh* (glyceraldehyde-3-phosphate dehydrogenase). The oligonucleotides used in this study are provided in Table supplement 5.

### Bacterial killing assay

BMDMs were plated at 2.5×10^5^ cells/well in a 12-well plate and allowed to adhere overnight. The cells were then infected with *Salmonella* (MOI = 10) for 30 min by centrifuging the plate at 1,000 rpm for 10 min and incubating for an additional 20 min. The plate was then rinsed twice with PBS containing 100 μg/ml gentamicin (Sigma, #345814) to remove the remaining extracellular bacteria. The cells were then incubated in fresh medium containing 10 μg/ml gentamicin to prevent the growth of extracellular bacteria. Where indicated, infected cells were lysed with 0.1% Triton X-100, and serial dilutions of the cell lysates were plated on TSA agar followed by incubation at 37°C overnight. The number of colonies appearing on the agar plate were counted in order quantify the surviving bacteria. Intracellular killing of macrophages was verified after 24-hour infection using transmission electron microscopy (TEM). BMDMs were harvested and subjected to routine processing, post-fixed, embedded, sectioned, and mounted at the Center for Cryo-Electron Microscopy (CCEM), Zhejiang University. Finally, thin sections (∼70 nm) were examined in an FEI Tecnai 10 (100 kV) transmission electron microscope.

### Nitric oxide (NO) measurement

Nitric oxide (NO) was determined through the formation of nitrite (NO_2_^-^), the primary, stable metabolite of NO. Extracellular nitrite released from BMDMs (1×10^5^ cells/well) stimulated with either HK-ST (MOI = 10) or 100 ng/ml LPS (Sigma, #L7895) was measured using a colorimetric assay based on the Griess reaction *(Green et al., 1982)*. At the indicated time points, cell culture supernatants were collected and placed into a fresh 96-well plate. The supernatants were then mixed with freshly prepared Griess solution consisting of 0.1% N-1-naphthylethylenediamine dihydrochloride (Sigma, #N9125) and 1% sulfanilamide (Sigma, #V900220) in 5% phosphoric acid (Sigma, #P5811) at a ratio of 1:1. The plate was incubated at room temperature for 10 min in the dark. Absorbance at 540 nm was then measured using a microplate reader (Biotek Eon, Gen5), and nitrite production was calculated using a NaNO_2_ calibration curve.

### Cell viability

Cell viability of BMDMs under high zinc conditions was measured using the plate-based colorimetric tetrazolium salt assay. First, BMDMs were plated in 96-well plates at 2×10^4^ cells/well. The next day, 2-fold serial dilutions of ZnSO_4_ (Sigma, #Z0251) were added to the culture media, and the cells were incubated at 37°C for 24 h. Next, the cell culture medium was carefully removed and replaced with 100 µl of fresh medium containing 5 mg/ml thiazolyl blue tetrazolium blue (Sigma, #M2128). The cells were incubated for an additional 4 h, and then the cell culture media was discarded. Blue crystals catalyzed by the mitochondrial enzyme succinate dehydrogenase were solubilized in 100% DMSO and the intensity was measured colorimetrically at 570 nm.

### Intracellular zinc measurement

A cell-based microplate assay was used to measure intracellular free zinc ions (Zn^2+^) using the cell-permeable fluorescent probe FluoZin-3 AM (Invitrogen, #F24195). BMDMs were plated into 96-well clear-bottom black polystyrene microplates (Corning, #3603) at 1×10^5^ cells/well and allowed to attach onto the bottom surface overnight. The cells were then incubated in culture media containing 2 μM FluoZin-3 AM for 30 min in the dark. After washing twice with PBS, the cells were incubated in fresh PBS for an additional 10 min to allow de-esterification. To detect cellular zinc accumulation, 100 μM ZnSO_4_ (Sigma, #Z0251) and/or 4 μM *N*,*N*,*N’*,*N’*-tetrakis-(2-pyridyl-methyl)-ethylenediamine (TPEN; Sigma, #P4413) was added to the cells, and FluoZin-3 fluorescence was measured at one-minute intervals for 30 min at 37°C using a SynergyMx M5 fluorescence microplate reader (Molecular Devices) with 485-nm excitation and 535-nm emission *(Brieger et al., 2013)*. The levels of intracellular zinc in response to pathogens and H_2_O_2_ were measured after 30-min stimulation. Images of FluoZin-3 fluorescence were acquired using a Nikon A1R confocal microscopy and analyzed using NIS-Elements Viewer imaging software, version 4.50.

### Intracellular ROS measurement

ROS production in BMDMs was measured using the membrane-permeable fluorescent probe 2’,7’-dichlorodihydrofluorescein diacetate (H_2_DCFHDA) (Sigma, #D6883). BMDMs were plated in 96-well plates at 1×10^5^ cells/well. The following morning, the cells were stained with 10 μM H_2_DCFHDA for 30 min at 37°C in the dark. After labeling, the cells were then washed twice with PBS, stimulated with HK-ST (MOI = 100) or LPS (1 µg/ml) for an additional 30 min at 37°C in the dark. The intensity of DCF fluorescence was then measured using a SpectraMax iD5 multi-mode microplate reader (Molecular Devices) with an excitation wavelength of 485 nm and an emission wavelength of 527 nm.

### Cell death assay

BMDMs were seeded in 6-well plates at 1×10^6^ cells/well and allowed to attach to the bottom surface overnight. The cells were then infected with *Salmonella* (MOI = 10) for 30 min, rinsed twice with PBS containing gentamicin (100 µg/ml), followed by perfused fresh culture media. At indicated time points, the cells were harvested and washed twice before subjecting to propidium iodide (PI) solution (10 µg/ml) (BioLegend, #421302) for 10 min, then the samples were mixed gently in the dark. Dead cells identified based on PI staining were measured using flow cytometry. When performing image analysis, BMDMs were seeded to glass-bottom dishes (Cellvis, #D35C4-20-1.5-N) and treated as described above. The cells were then gently washed and stained with PI in PBS containing 2% FBS for 10 min, and imaged using an Olympus CSU-W1 confocal microscope.

### Protein quantification using western blot analysis

Proteins from BMDMs were extracted using RIPA buffer (Solarbio, #R0020) containing Pierce protease inhibitor (Thermo Fisher Scientific, #A32965) and phosphatase inhibitor (Roche, #04906845001). Total protein content was quantified using the Bradford dye-binding method (Sigma, #B6916). A total of 50 μg of cellular proteins were mixed with 5x loading buffer and boiled for 5 min. Equal amounts of prepared proteins were electrophoresed in 10% SDS-PAGE gel at 100 V. After gel electrophoresis, separated proteins were transferred onto polyvinylidene difluoride (PVDF) membrane (Bio-Rad, #1620177) at 300 mA for 90 min using transfer buffer containing 25 mM Tris, 192 mM glycine, and 20% methanol. The membranes were then blocked in 5% (w/v) skim milk in Tris-buffered saline containing 0.05% Tween-20 (TBST) for 90 min at room temperature. The membranes were then washed three times with TBST for 10 min each, and then probed using the following primary antibodies overnight at 4°C (1:1000 dilution): iNOS antibody (#39898S), NF-κB p65 (D14E12) XP rabbit mAb (#8242S), or phospho-NF-κB p65 (Ser536) (93H1) rabbit mAb (#3033) (all from Cell Signaling Technology). A monoclonal anti-FLAG M2-Peroxidase (HRP) antibody was used to detect the flag-tagged fusion protein. The following day, the membranes were washed with TBST and incubated with a secondary antibody, HRP-conjugated goat anti-rabbit IgG (H+L) (ABclonal, #AS014) diluted in 5% milk/TBST (1:2000) at room temperature for 2 h. The membranes were then washed in TBST three times for 10 min each to remove excessive antibody and developed using Pierce^TM^ ECL Western Blotting Substrate (Thermo Fisher Scientific, #32106). The bands were quantified using the ChemiDoc Touch Imaging System (Bio-Rad).

### Statistical analysis

Statistical significance between two groups was determined using a 2-tailed, unpaired Student *t*-test (for comparing two groups). Data are presented as mean ± SEM. For comparison of survival curves, Log-rank (Mantel-Cox) test was performed. The sample size for each statistical analysis is provided in the figure legends. Data were analyzed and generated using GraphPad Prism version 7.04 (GraphPad Software Inc., La Jolla, CA, USA). For all studies, *P* values below 0.05 was considered to be statistically significant. \**P*<0.05; \*\**P*<0.01; \*\*\**P*<0.001; ns, not significant.

## Acknowledgments

This work was supported by the National Natural Science Foundation of China (31930057 to F.W. and 31970689 to J.M.).

## Funding

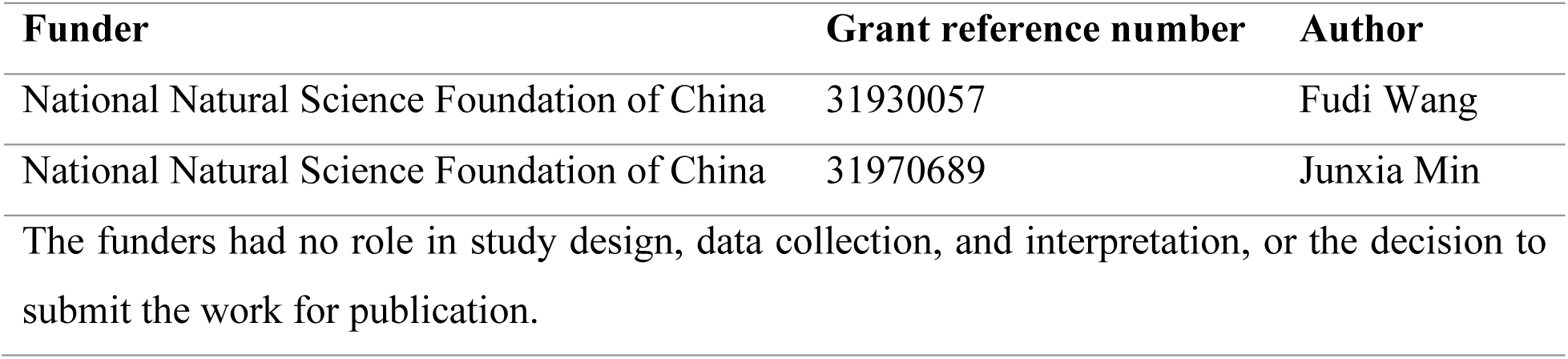

## Author contributions

Pinanong Na-Phatthalung, Junxia Min, and Fudi Wang designed the study; Pinanong Na-Phatthalung, Shumin Sun, and Enjun Xie performed the research and analyzed the data; Pinanong Na-Phatthalung, Shumin Sun, Enjun Xie, and Jia Wang contributed new reagents and/or analytic tools; and Pinanong Na-Phatthalung, Junxia Min, and Fudi Wang wrote the paper.

## Author ORCIDs

Fudi Wang https://orcid.org/0000-0001-8730-0003

Junxia Min https://orcid.org/0000-0001-8099-6327

Pinanong Na-Phatthalung https://orcid.org/0000-0002-8162-4645

## Ethics

Animal experimentation: All *in vivo* experiments were conducted in strict accordance with the recommendations in the Guide for the Care and Use of Laboratory Animals of the National Institutes of Health. All animal experiments were approved by the Institutional Animal Care and Use Committee, Zhejiang University.

## Additional files

### Supplementary files

Supplementary file 1

## Data availability

Figure 1 - Source data 1 contains the RNA-seq datasets used to generate figure 1F and the data have been deposited in NCBI Gene Expression Omnibus ID GSE67427.

## Supplementary Figures

**Figure supplement 1.**
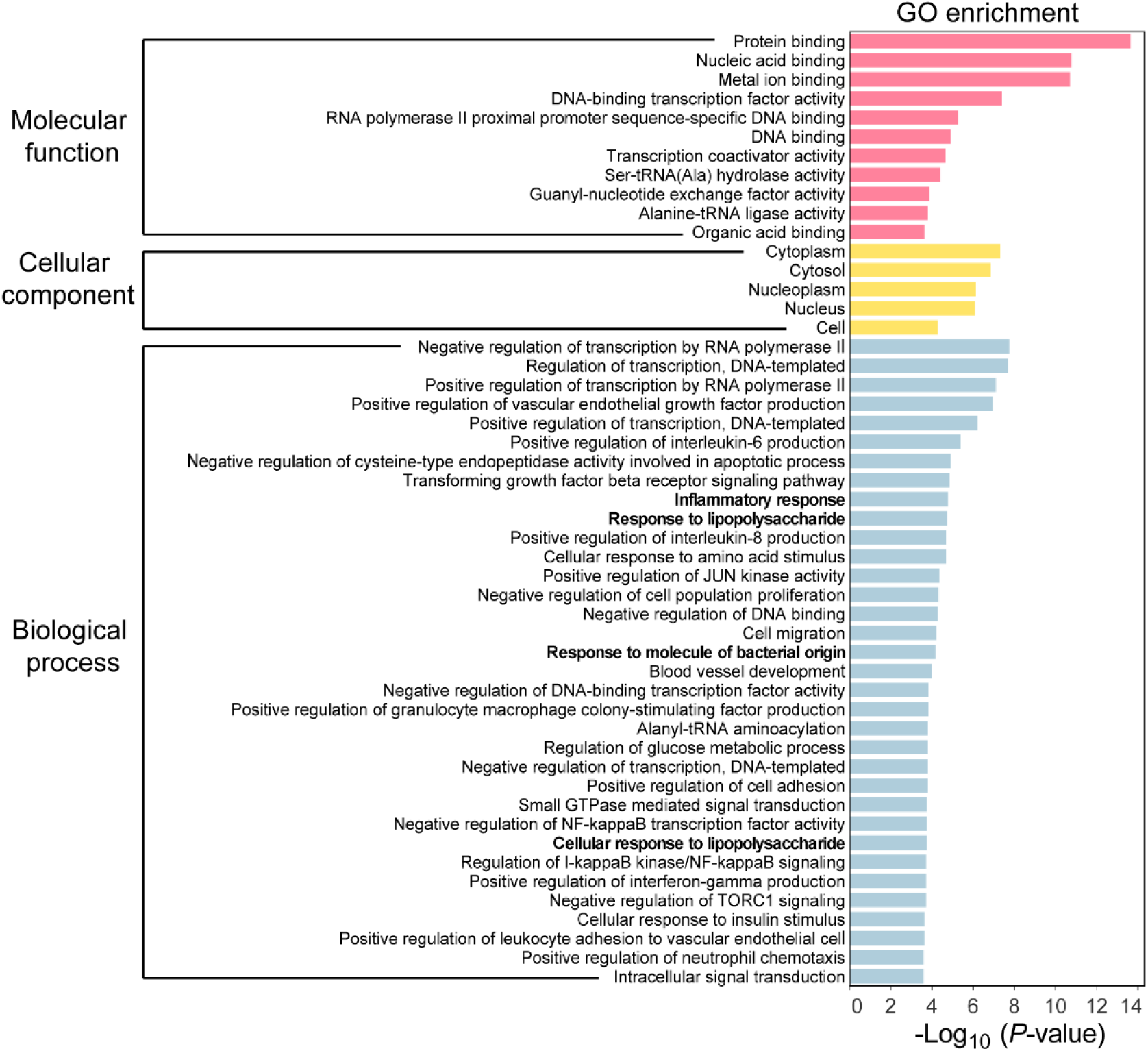
Gene ontology (GO) enrichment analysis of differentially expressed genes (DEGs) in BMDMs with or without *Salmonella* infection. The bar graph shows the top 50 GO terms categorized by molecular function, cellular component, and biological process in BMDMs infected with Salmonella (MOI = 1) for 2 h compared to uninfected BMDMs.

**Figure supplement 2.**
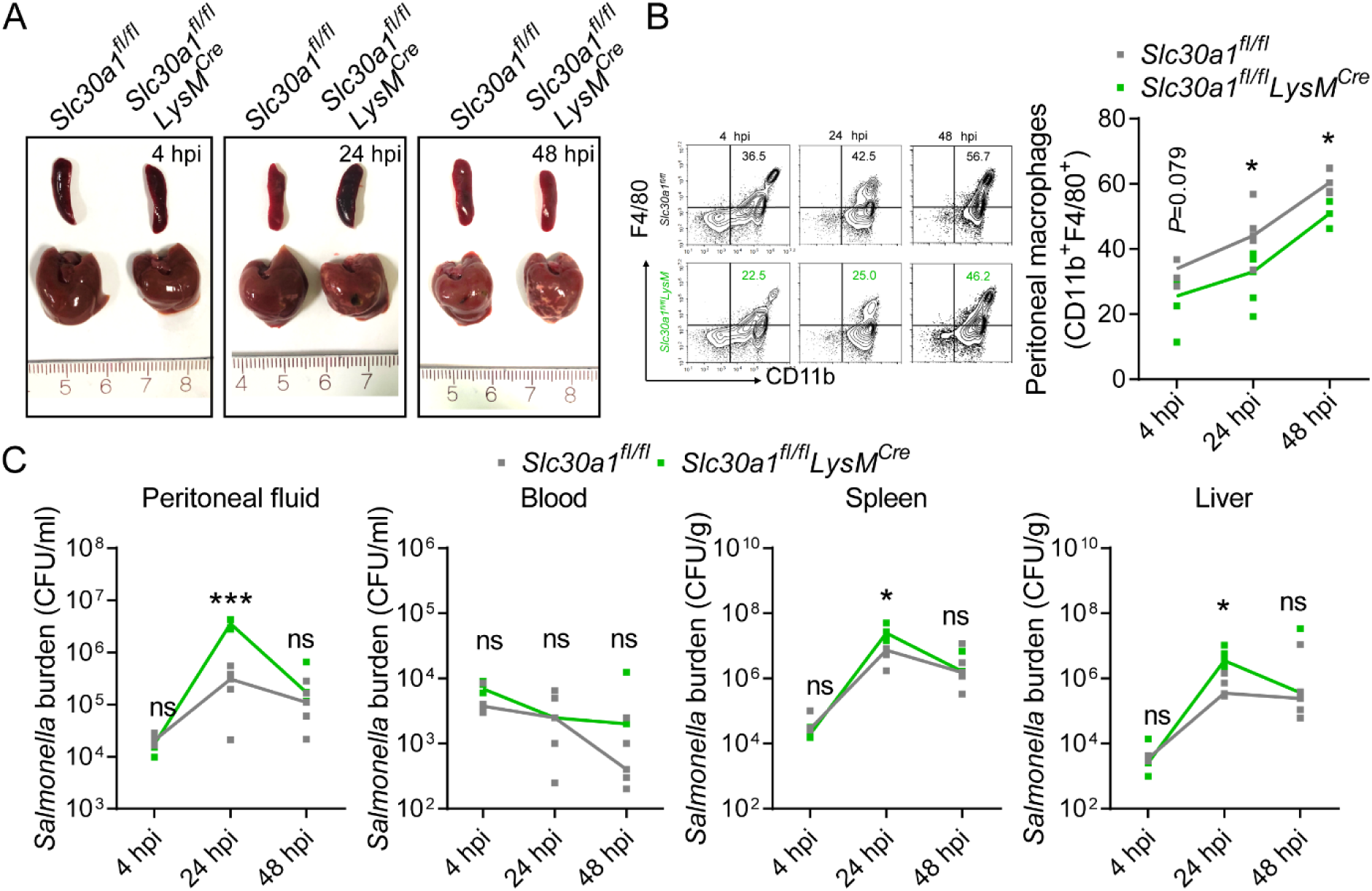
*Salmonella* infection leads to significantly decreased macrophages and increased tissue bacterial burden in *Slc30a1^fl/fl^LysM^Cre^* mice. (**A**) Gross image of spleen and liver of mice after infection with a non-lethal dose of *Salmonella* (1×10^4^ CFU/mouse) 4, 24, and 48 hpi. (**B**) FACS plots of CD11b^+^F4/80^+^ peritoneal macrophages obtained from *Salmonella*-infected *Slc30a1^fl/fl^* and *Slc30a1^fl/fl^LysM^Cre^* mice (n = 3-5 mice/group). (**C**) Bacterial burden (in CFU/ml) measured in the peritoneal fluid, blood, spleen, and liver of *Salmonella*-infected *Slc30a1^fl/fl^* and *Slc30a1^fl/fl^LysM^Cre^* mice (n = 3-5 mice/group). Data in this figure are represented as mean ± SEM. *P* values were determined using 2-tailed unpaired Student’s *t*-test. \**P*<0.05, \*\**P*<0.01, \*\*\**P*<0.001 and ns, not significant.

**Figure supplement 3.**
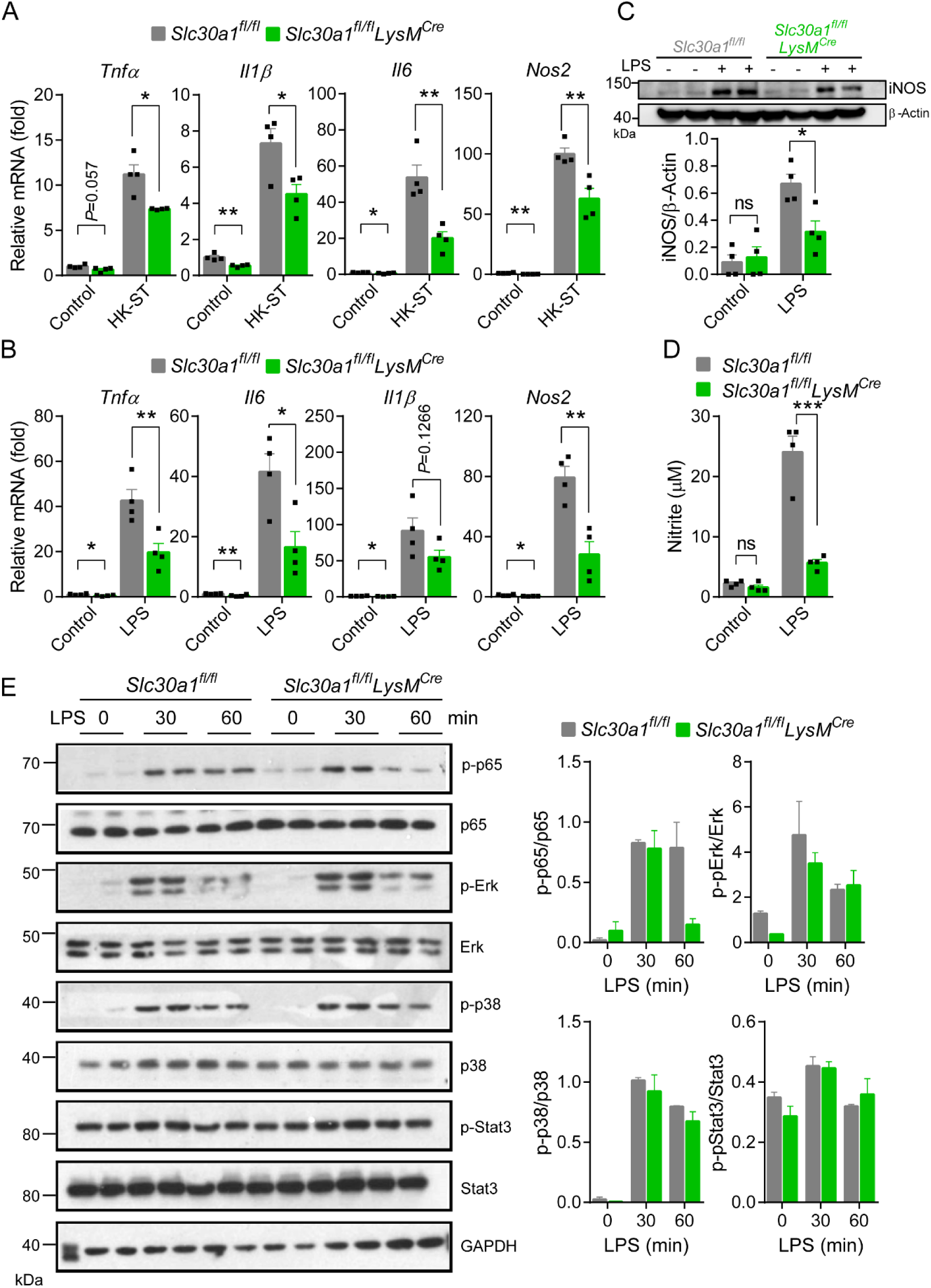
Loss of *Slc30a1* in the murine macrophages significantly reduces iNOS and NO production by inhibiting the NF-κB pathway. (**A** and **B**) Summary of *Tnfα*, *Ilβ*, *Il6*, and *Nos2* mRNA measured in *Slc30a1^fl/fl^* and *Slc30a1^fl/fl^LysM^Cre^* BMDMs after 4 h of HK-ST (MOI = 10) or LPS (100 ng/ml) stimulation relative to unstimulated cells (*n* = 4). (**C**) Western blot analysis and summary of iNOS protein in BMDMs after LPS stimulation (100 ng/ml) for 24 h (*n* = 4). (**D**) Summary of nitrite concentration measured in the cell culture supernatant of BMDMs after LPS treatment for 24 h (*n* = 4). (**E**) Western blot analysis and summary of p-p65, p65, p-Erk, Erk, p-p38, p38, p-Stat3, and Stat3 measured in BMDMs after LPS treatment (100 ng/ml) for 0, 30, and 60 min (*n* = 2). Data in this figure are represented as mean ± SEM. *P* values were determined using 2-tailed unpaired Student’s *t*-test. \**P*<0.05, \*\**P*<0.01, \*\*\**P*<0.001 and ns, not significant.

**Figure supplement 4.**
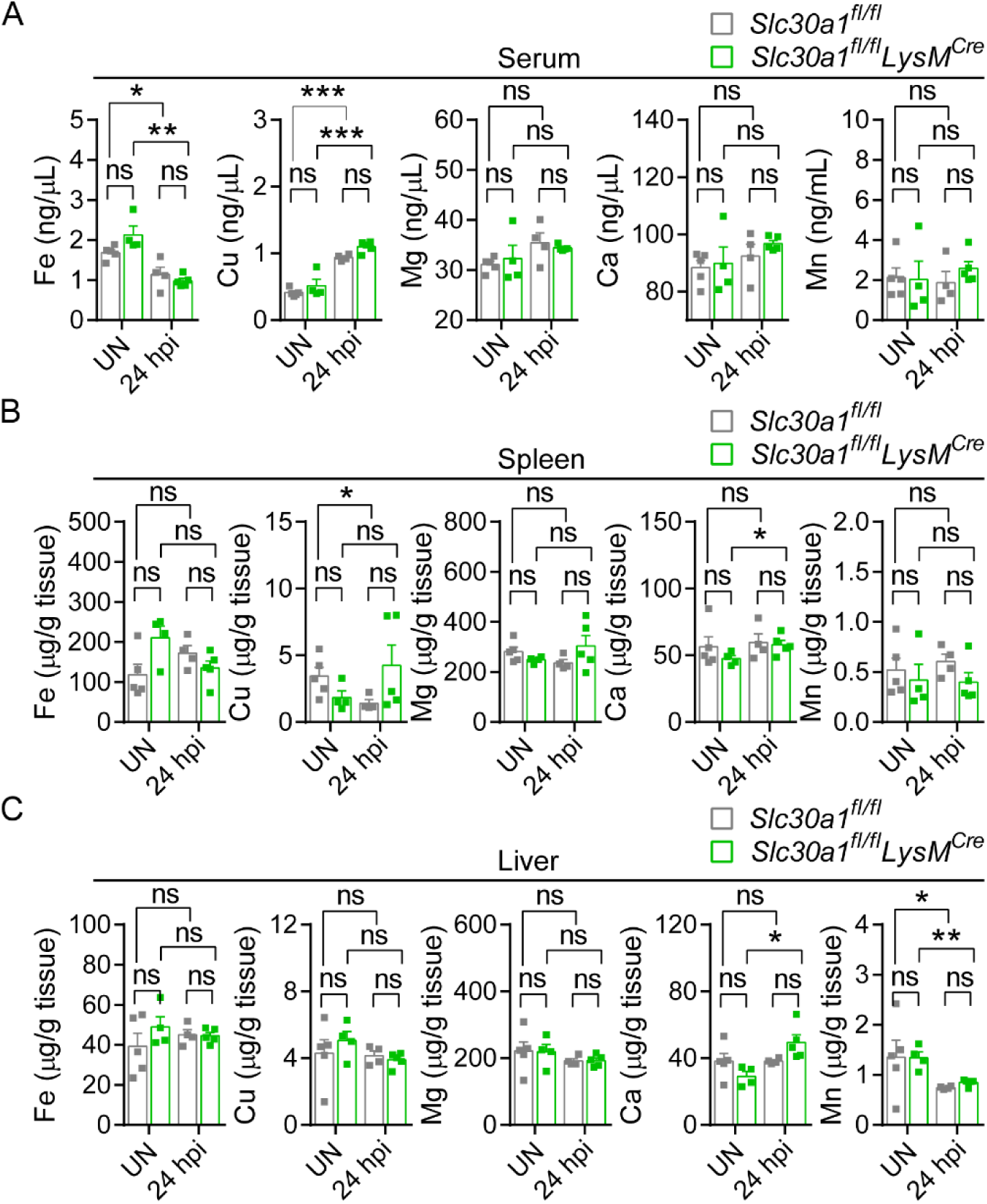
Except for Zn, macrophage specific deletion of *Slc30a1* in mice does not affect the content of other trace minerals. **(A-C)** Summary of Fe, Cu, Mg, Ca, and Mn content measured in the serum (**A**), spleen (**B**), and liver (**C**) of uninfected (UN) and *Salmonella*-infected mice at 24 hpi (*n* = 4-5 mice/group). Data in this figure are represented as mean ± SEM. *P* values were determined using 2-tailed unpaired Student’s *t*-test. \**P*<0.05, \*\**P*<0.01, \*\*\**P*<0.001 and ns, not significant.

**Figure supplement 5.**
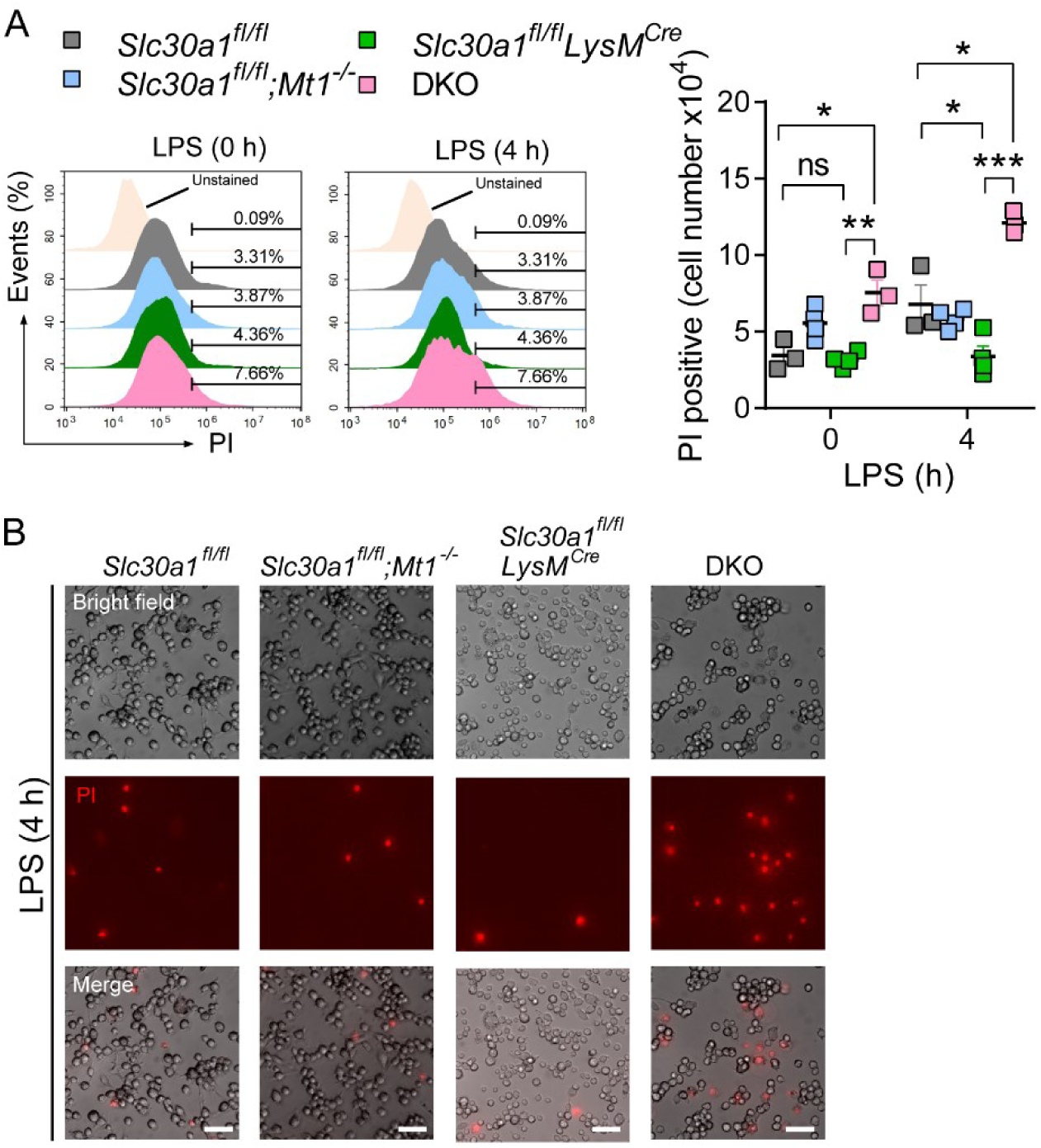
Loss of both *Slc30a1* and *Mt1* increases LPS-induced cell death of macrophages. (**A**) FACS analysis and summary of propidium iodide (PI)‒positive BMDMs either untreated for treated with LPS (1 µg/ml) for 4 h (*n* = 3-4). (**B**) Bright-field and fluorescence microscopy images of PI-stained BMDMs after LPS treatment for 4 h. Scale bars, 50 µm. Data in this figure are represented as mean ± SEM. *P* values in **A** were determined using 2-tailed unpaired Student’s *t*-test. \**P*<0.05, \*\**P*<0.01, \*\*\**P*<0.001 and ns, not significant.

**Figure supplement 6.**
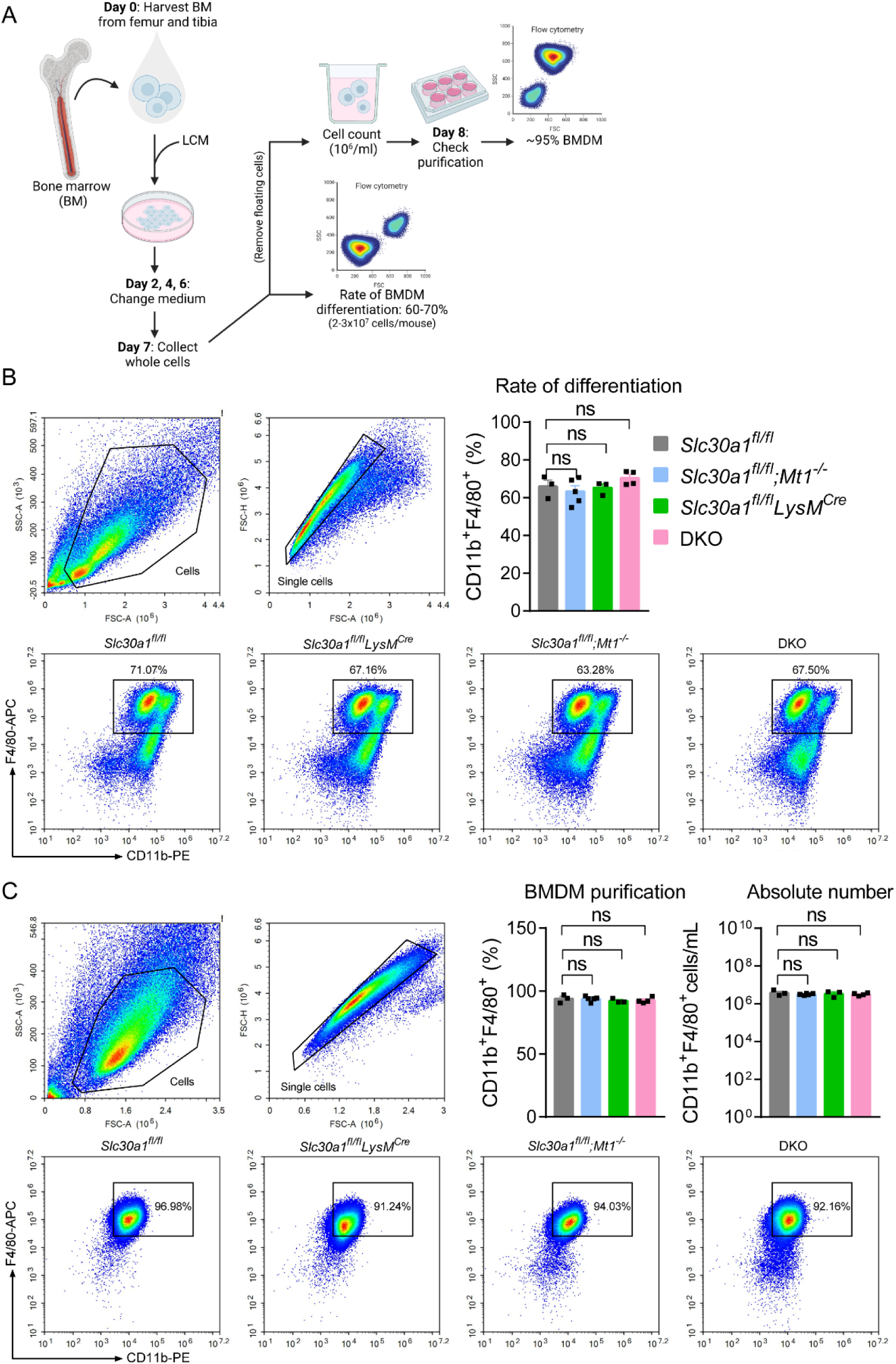
Macrophage specific deletion of *Slc30a1* in mice does not affect the differentiation of BMDMs. **(A)** Schematic illustration depicting the strategy used to measure the BMDM differentiation procedure. Bone marrow (BM) cells were harvested from the femur and tibia and cultured in L929 cell-conditioned medium (LCM) for 7 days. (**B**) Flow cytometry analysis and summary of the percent BMDM differentiation (CD11b^+^F4/80^+^) from BM samples obtained from the indicated mice measured at day 7 (*n* = 3-5). (**C**) Percent BMDM purification and absolute number of BMDMs measured after removing floating cells and measuring cell concentration. Data in this figure are represented as mean ± SEM. *P* values were determined using 2-tailed unpaired Student’s *t*-test. ns, not significant.

## Supplementary tables

**Table supplement 1.** Top 50 enriched GO terms for the DEGs in C57BL/6 BMDMs infected with *Salmonella* versus uninfected cells.

|  | GO ID | Term | Function | P-value |
| --- | --- | --- | --- | --- |
| 1 | GO:0005515 | Protein binding | Molecular function | 2.31E-14 |
| 2 | GO:0003676 | Nucleic acid binding | Molecular function | 1.65E-11 |
| 3 | GO:0046872 | Metal ion binding | Molecular function | 2.02E-11 |
| 4 | GO:0000122 | Negative regulation of transcription by RNA polymerase II | Biological process | 1.77E-08 |
| 5 | GO:0006355 | Regulation of transcription, DNA-templated | Biological process | 2.08E-08 |
| 6 | GO:0003700 | DNA-binding transcription factor activity | Molecular function | 4.02E-08 |
| 7 | GO:0005737 | Cytoplasm | Cellular component | 5.18E-08 |
| 8 | GO:0045944 | Positive regulation of transcription by RNA polymerase II | Biological process | 7.75E-08 |
| 9 | GO:0010575 | Positive regulation of vascular endothelial growth factor production | Biological process | 1.12E-07 |
| 10 | GO:0005829 | Cytosol | Cellular component | 1.36E-07 |
| 11 | GO:0045893 | Positive regulation of transcription, DNA-templated | Biological process | 6.28E-07 |
| 12 | GO:0005654 | Nucleoplasm | Cellular component | 7.75E-07 |
| 13 | GO:0005634 | Nucleus | Cellular component | 8.66E-07 |
| 14 | GO:0032755 | Positive regulation of interleukin-6 production | Biological process | 4.31E-06 |
| 15 | GO:0000978 | RNA polymerase II proximal promoter sequence-specific DNA binding | Molecular function | 5.41E-06 |
| 16 | GO:0003677 | DNA binding | Molecular function | 1.27E-05 |
| 17 | GO:0043154 | Negative regulation of cysteine-type endopeptidase activity involved in apoptotic process | Biological process | 1.27E-05 |
| 18 | GO:0007179 | Transforming growth factor beta receptor signaling pathway | Biological process | 1.45E-05 |
| 19 | GO:0006954 | **Inflammatory response** | Biological process | 1.73E-05 |
| 20 | GO:0032496 | **Response to lipopolysaccharide** | Biological process | 1.89E-05 |
| 21 | GO:0032757 | Positive regulation of interleukin-8 production | Biological process | 1.97E-05 |
| 22 | GO:0071230 | Cellular response to amino acid stimulus | Biological process | 2.06E-05 |
| 23 | GO:0003713 | Transcription coactivator activity | Molecular function | 2.17E-05 |
| 24 | GO:0002196 | Ser-tRNA(Ala) hydrolase activity | Molecular function | 3.97E-05 |
| 25 | GO:0043507 | Positive regulation of JUN kinase activity | Biological process | 4.30E-05 |
| 26 | GO:0008285 | Negative regulation of cell population proliferation | Biological process | 4.94E-05 |
| 27 | GO:0005623 | Cell | Cellular component | 5.17E-05 |
| 28 | GO:0043392 | Negative regulation of DNA binding | Biological process | 5.41E-05 |
| 29 | GO:0016477 | Cell migration | Biological process | 6.52E-05 |
| 30 | GO:0002237 | **Response to molecule of bacterial origin** | Biological process | 7.11E-05 |
| 31 | GO:0001568 | Blood vessel development | Biological process | 9.90E-05 |
| 32 | GO:0005085 | Guanyl-nucleotide exchange factor activity | Molecular function | 1.34E-04 |
| 33 | GO:0043433 | Negative regulation of DNA-binding transcription factor activity | Biological process | 1.43E-04 |
| 34 | GO:0032725 | Positive regulation of granulocyte macrophage colony-stimulating factor production | Biological process | 1.48E-04 |
| 35 | GO:0006419 | Alanyl-tRNA aminoacylation | Biological process | 1.55E-04 |
| 36 | GO:0004813 | Alanine-tRNA ligase activity | Molecular function | 1.55E-04 |
| 37 | GO:0010906 | Regulation of glucose metabolic process | Biological process | 1.58E-04 |
| 38 | GO:0045892 | Negative regulation of transcription, DNA-templated | Biological process | 1.62E-04 |
| 39 | GO:0045785 | Positive regulation of cell adhesion | Biological process | 1.65E-04 |
| 40 | GO:0007264 | Small GTPase mediated signal transduction | Biological process | 1.71E-04 |
| 41 | GO:0032088 | Negative regulation of NF-kappaB transcription factor activity | Biological process | 1.81E-04 |
| 42 | GO:0071222 | **Cellular response to lipopolysaccharide** | Biological process | 1.84E-04 |
| 43 | GO:0043122 | Regulation of I-kappaB kinase/NF-kappaB signaling | Biological process | 2.00E-04 |
| 44 | GO:0032729 | Positive regulation of interferon-gamma production | Biological process | 2.01E-04 |
| 45 | GO:1904262 | Negative regulation of TORC1 signaling | Biological process | 2.02E-04 |
| 46 | GO:0032869 | Cellular response to insulin stimulus | Biological process | 2.26E-04 |
| 47 | GO:0043177 | Organic acid binding | Molecular function | 2.41E-04 |
| 48 | GO:1904996 | Positive regulation of leukocyte adhesion to vascular endothelial cell | Biological process | 2.41E-04 |
| 49 | GO:0090023 | Positive regulation of neutrophil chemotaxis | Biological process | 2.50E-04 |
| 50 | GO:0035556 | Intracellular signal transduction | Biological process | 2.68E-04 |

**Table supplement 2.**
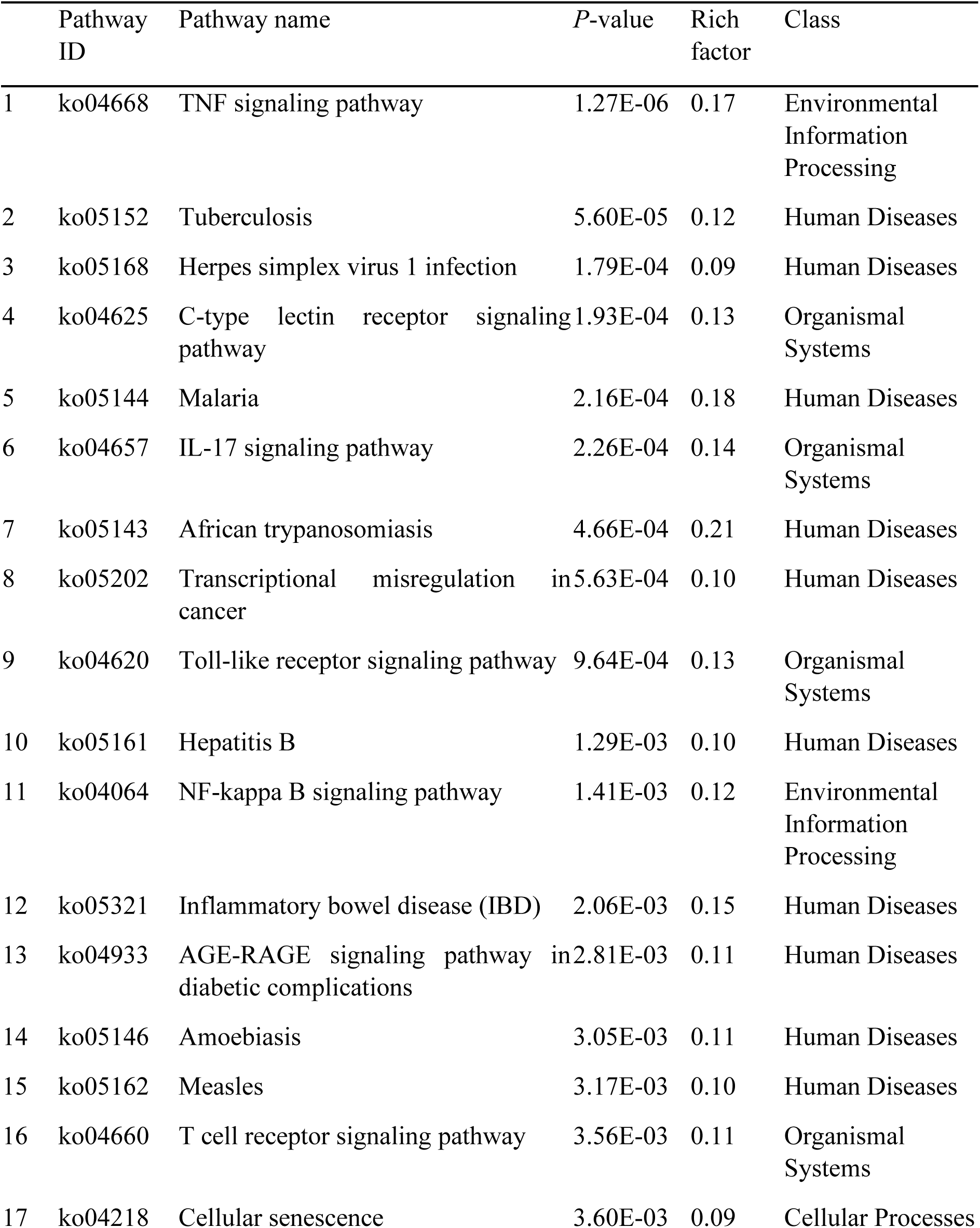
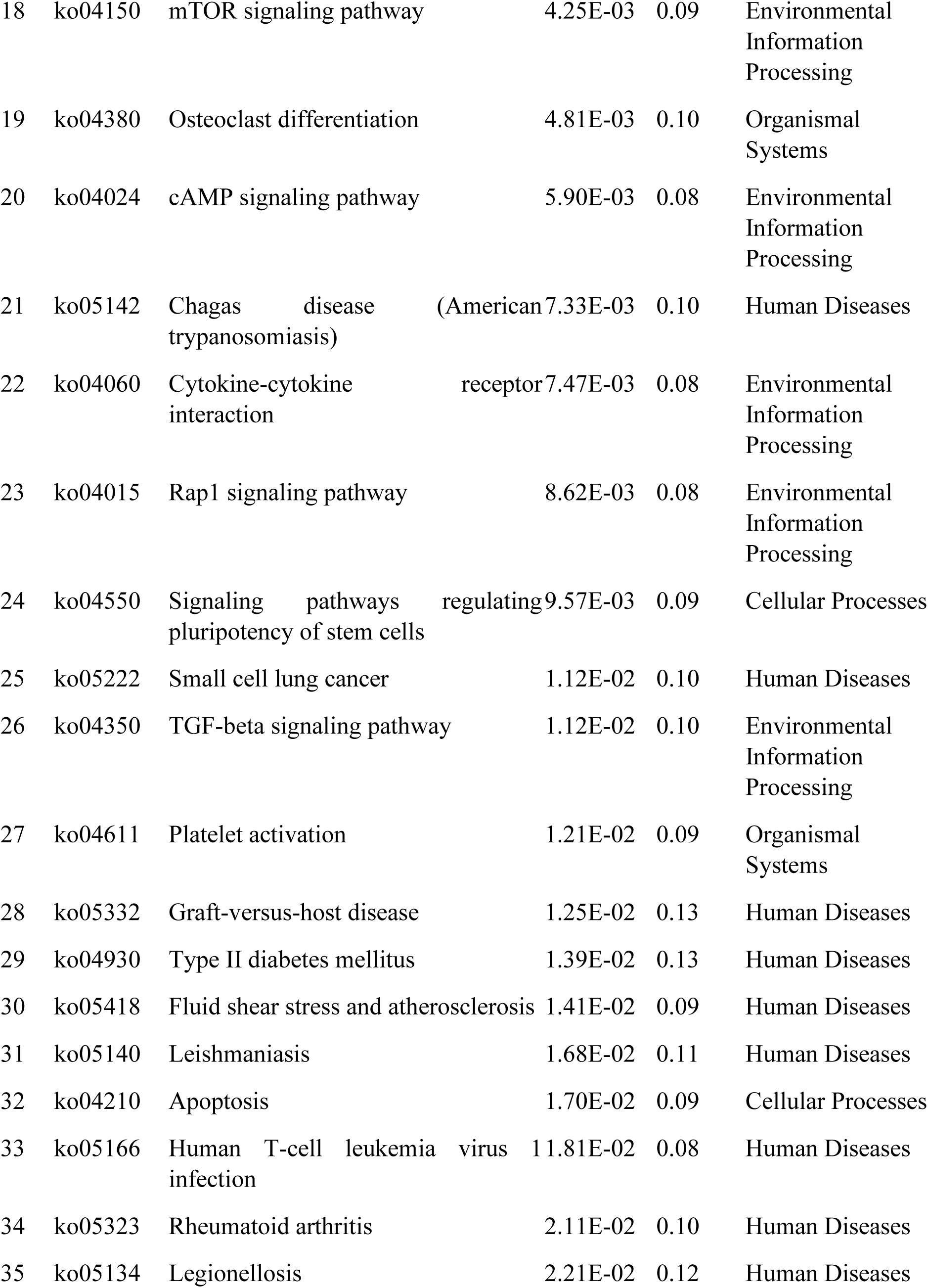
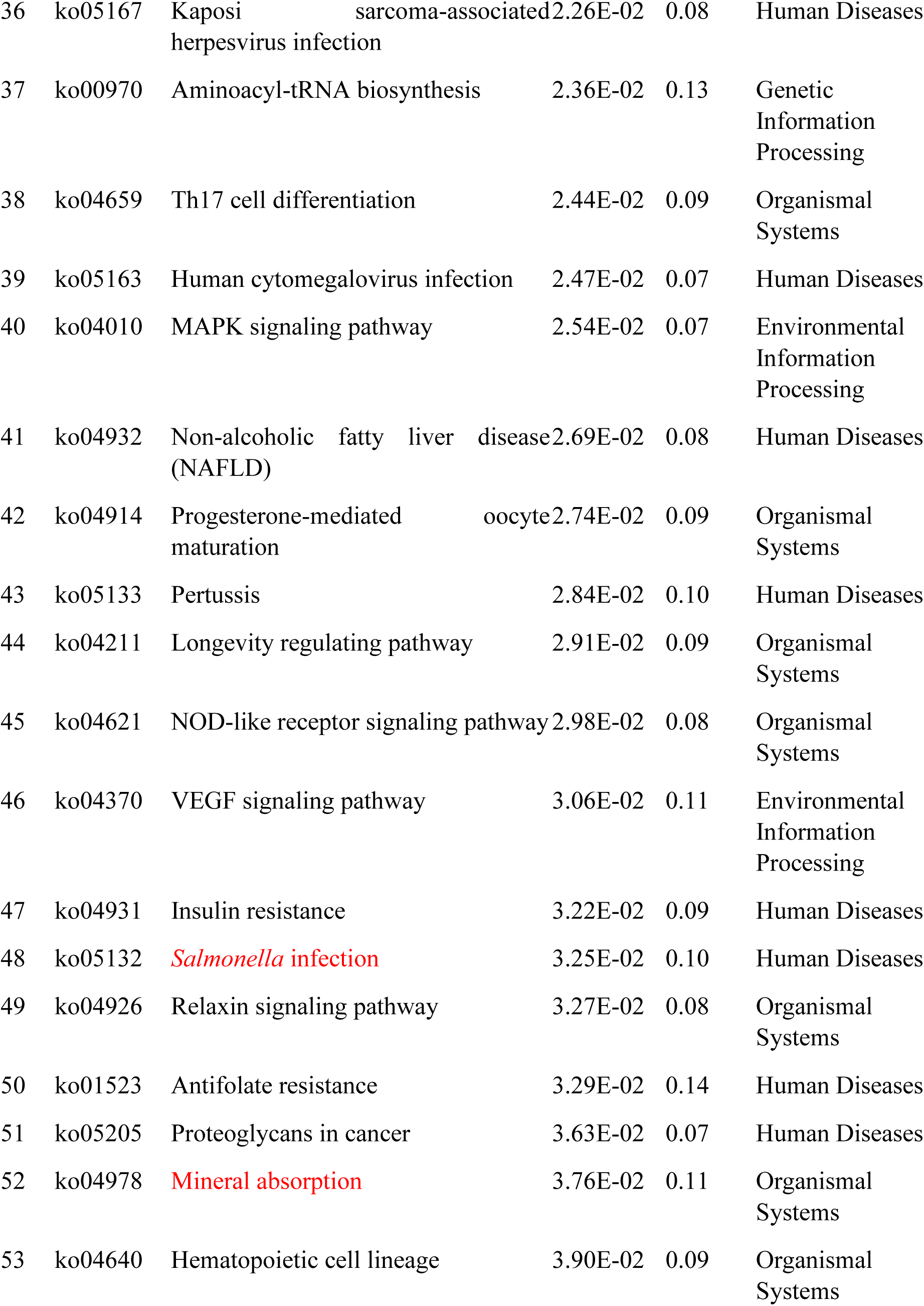
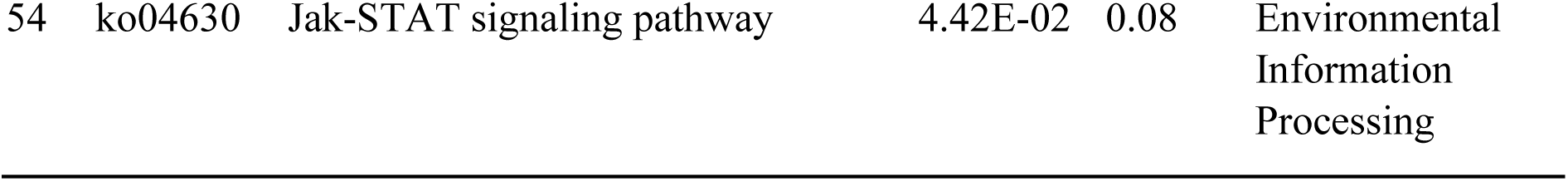
KEGG enrichment pathways of the DEGs in C57BL/6 BMDMs infected with *Salmonella* versus uninfected cells.

**Table supplement 3.** Summary of blood test results for uninfected and *Salmonella*-infected *Slc30a1^fl/fl^*and *Slc30a1^fl/fl^LysM^Cre^* mice.

| Measurement | *Slc30a1<sup>fl/fl</sup>* | *Slc30a1<sup>fl/fl</sup>LysM<sup>Cre</sup>* | *Slc30a1<sup>fl/fl</sup>* | *Slc30a1<sup>fl/fl</sup>LysM<sup>Cre</sup>* |
| --- | --- | --- | --- | --- |
|  | Uninfected (*n* = 5) | Uninfected (*n* = 6) | 24 hpi (*n* = 5) | 24 hpi (*n* = 4) |
| WBC ($\times 10^9/L$) | 2.29±0.57 | 1.89±0.33 | 2.46±1.01 | 1.38±0.53 |
| RBC ($\times 10^{12}/L$) | 9.02±0.29 | 8.85±0.57 | 9.07±0.63 | 8.72±0.36 |
| Hb (g/dL) | 13.46±0.34 | 13.37±0.70 | 12.82±1.10 | 12.53±0.44 |
| Hct (%) | 43.56±1.45 | 43.42±2.19 | 40.98±2.99 | 40.68±1.19 |
| MCV (fL) | 48.26±0.84 | 49.08±1.15 | 45.22±1.96 | 46.63±0.59 |
| MCH (pg) | 14.94±0.25 | 15.12±0.22 | 14.14±0.52 | 14.35±0.06 |
| MCHC (g/L) | 30.90±0.41 | 30.80±0.52 | 31.26±0.67 | 30.80±0.22 |
| PLT ($\times 10^9/L$) | 1301 ±111 | 1086±210 | 717±99 | 788±148 |
| LYM ($\times 10^9/L$) | 1.61±0.42 | 1.44±0.09 | 1.45±0.40 | 0.77±0.25 |
| EO ($\times 10^9/L$) | 0.04±0.01 | 0.02±0.02 | 0.04±0.03 | 0.02±0.01 |
| BASO ($\times 10^9/L$) | 0.00±0.01 | 0.00±0.00 | 0.05±0.11 | 0.02±0.03 |

hpi, hours post-infection; WBC, white blood cells; RBC, red blood cells; Hb, hemoglobin; Hct; hematocrit; MCV, mean corpuscular volume; MCH, mean corpuscular hemoglobin; MCHC, mean corpuscular hemoglobin concentration; PLT, platelets; LYM, lymphocytes; EO, eosinophils; BASO, basophils. Data are presented as the mean ± SEM.

**Table supplement 4.**
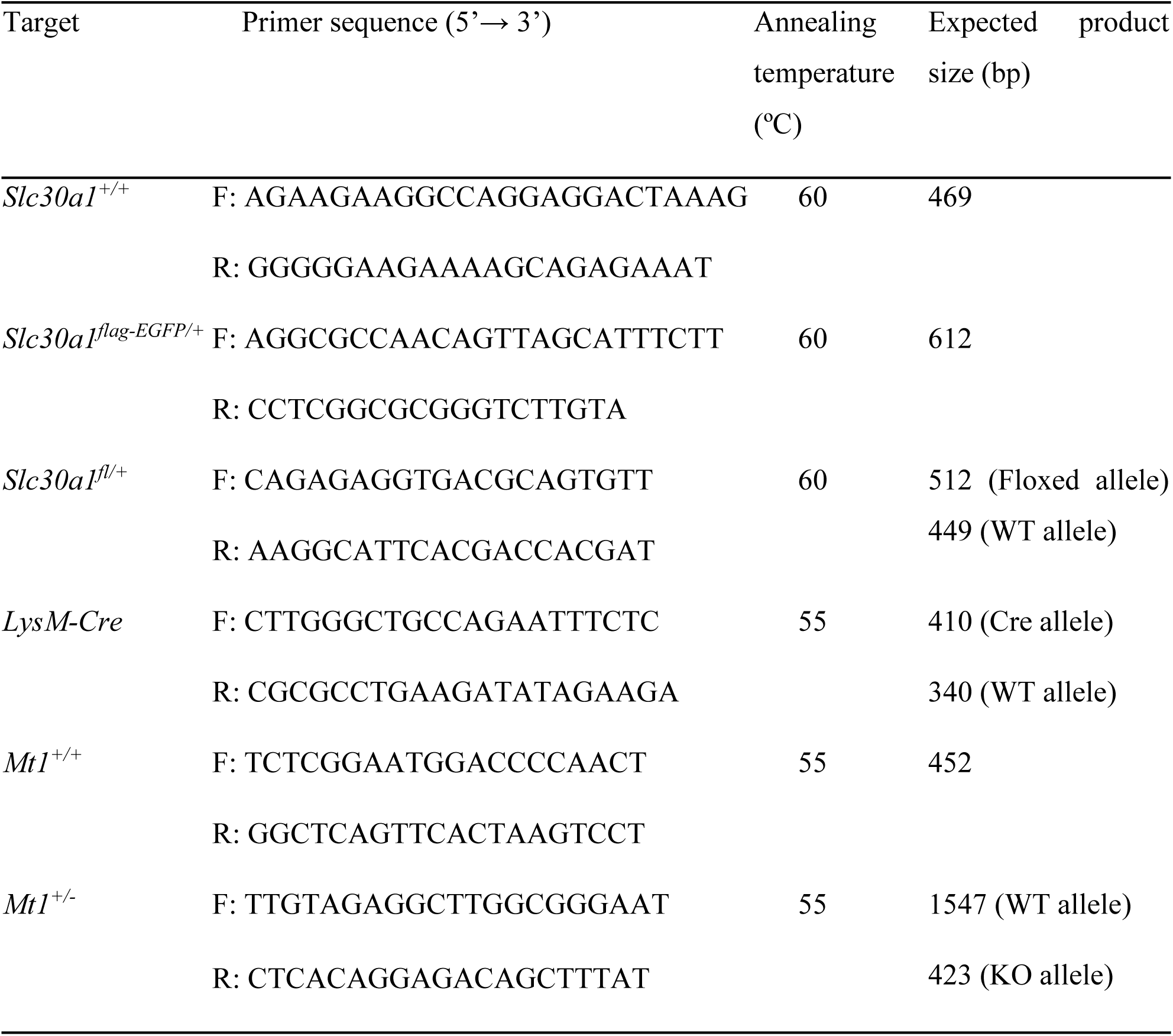
Primer pairs used for genotyping.

**Table supplement 5.**
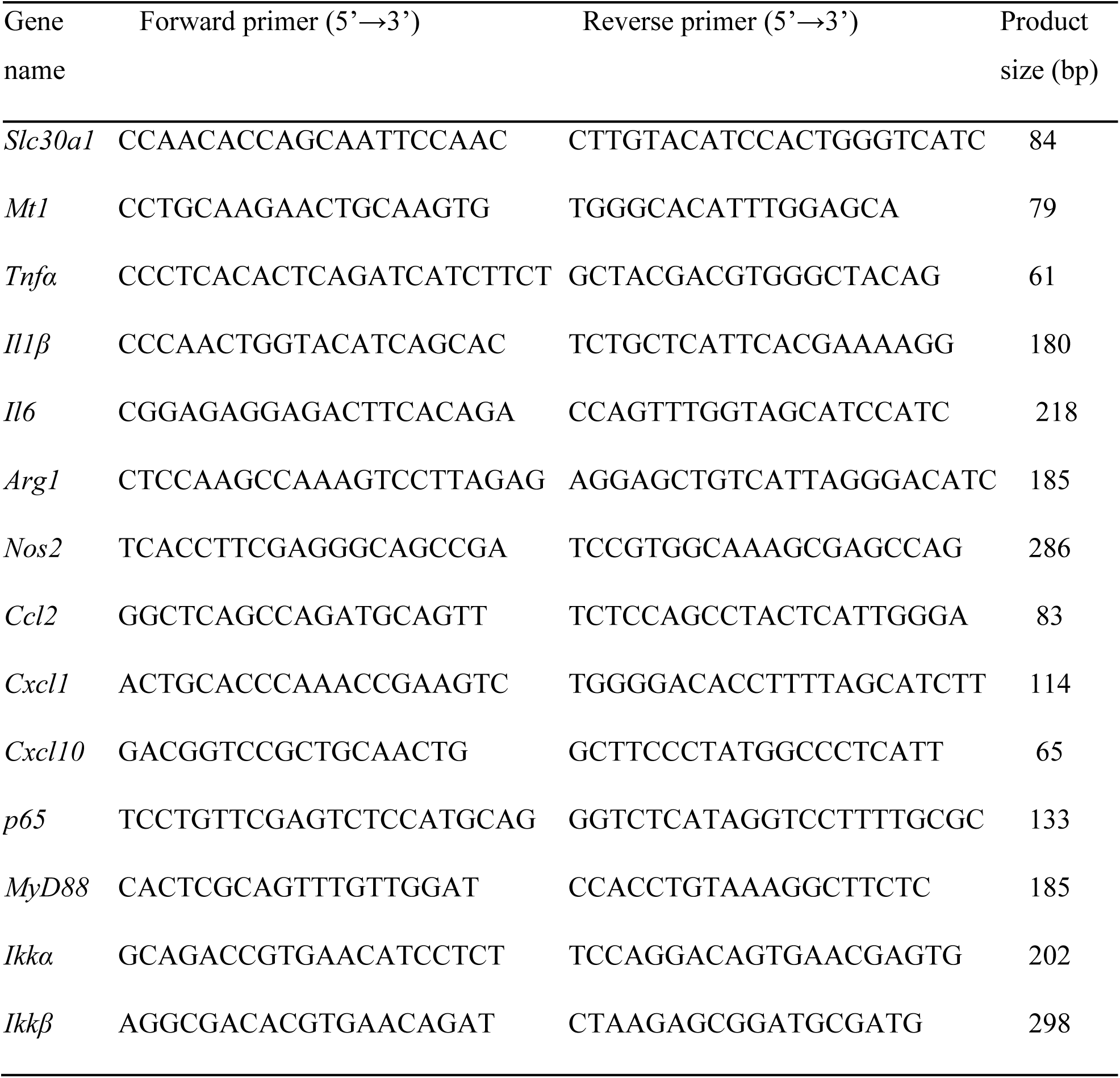
List of primer pairs used for RT-qPCR analysis.

## References

Ahuja A, Kim E, Sung GH, Cho JY. 2020. STAT3 differentially regulates TLR4-mediated inflammatory responses in early or late phases. International journal of molecular sciences 21: 7675. https://doi.org/10.3390/ijms21207675.

Alker W, Haase H. 2018. Zinc and sepsis. Nutrients 10. https://doi.org/10.3390/nu10080976.

Andrews GK, Wang H, Dey SK, Palmiter RD. 2004. Mouse zinc transporter 1 gene provides an essential function during early embryonic development. Genesis (New York, N.Y. : 2000) 40: 74-81. https://doi.org/10.1002/gene.20067.

Boone DL, Turer EE, Lee EG, Ahmad RC, Wheeler MT, Tsui C, Hurley P, Chien M, Chai S, Hitotsumatsu O, McNally E, Pickart C, Ma A. 2004. The ubiquitin-modifying enzyme A20 is required for termination of Toll-like receptor responses. Nature immunology 5: 1052–1060. https://doi.org/10.1038/ni1110.

Botella H, Peyron P, Levillain F, Poincloux R, Poquet Y, Brandli I, Wang C, Tailleux L, Tilleul S, Charrière GM, Waddell SJ, Foti M, Lugo-Villarino G, Gao Q, Maridonneau-Parini I, Butcher PD, Castagnoli PR, Gicquel B, de Chastellier C, Neyrolles O. 2011. Mycobacterial p(1)-type ATPases mediate resistance to zinc poisoning in human macrophages. Cell host & microbe 10: 248–259. https://doi.org/10.1016/j.chom.2011.08.006.

Brennan MA, Cookson BT. 2000. *Salmonella* induces macrophage death by caspase-1-dependent necrosis. Molecular microbiology 38: 31–40. https://doi.org/10.1046/j.1365-2958.2000.02103.x.

Brieger A, Rink L, Haase H. 2013. Differential regulation of TLR-dependent MyD88 and TRIF signaling pathways by free zinc ions. Journal of immunology (Baltimore, Md. : 1950) 191: 1808-1817. https://doi.org/10.4049/jimmunol.1301261.

Carreno D, Wanford JJ, Jasiunaite Z, Hames RG, Chung WY, Dennison AR, Straatman K, Martinez-Pomares L, Pareek M, Orihuela CJ, Restrepo MI, Lim WS, Andrew PW, Moxon ER, Oggioni MR. 2021. Splenic macrophages as the source of bacteraemia during pneumococcal pneumonia. EBioMedicine 72: 103601. https://doi.org/10.1016/j.ebiom.2021.103601.

Chan ED, Riches DW. 2001. IFN-gamma + LPS induction of iNOS is modulated by ERK, JNK/SAPK, and p38(mapk) in a mouse macrophage cell line. American journal of physiology. Cell physiology 280: C441-450. https://doi.org/10.1152/ajpcell.2001.280.3.C441.

Chandra RK. 1984. Excessive intake of zinc impairs immune responses. Jama 252: 1443–1446.

Clausen BE, Burkhardt C, Reith W, Renkawitz R, Förster I. 1999. Conditional gene targeting in macrophages and granulocytes using LysMcre mice. Transgenic research 8: 265–277. https://doi.org/10.1023/a:1008942828960.

Dai H, Wang L, Li L, Huang Z, Ye L. 2021. Metallothionein 1: a new spotlight on inflammatory diseases. Frontiers in immunology 12: 739918. https://doi.org/10.3389/fimmu.2021.739918.

Depaolo RW, Lathan R, Rollins BJ, Karpus WJ. 2005. The chemokine CCL2 is required for control of murine gastric *Salmonella enterica* infection. Infection and immunity 73: 6514–6522. https://doi.org/10.1128/iai.73.10.6514-6522.2005.

Dineley KE, Votyakova TV, Reynolds IJ. 2003. Zinc inhibition of cellular energy production: implications for mitochondria and neurodegeneration. Journal of neurochemistry 85: 563–570. https://doi.org/10.1046/j.1471-4159.2003.01678.x.

Du W, Gu M, Hu M, Pinchi P, Chen W, Ryan M, Nold T, Bannaga A, Xu H. 2021. Lysosomal Zn(2+) release triggers rapid, mitochondria-mediated, non-apoptotic cell death in metastatic melanoma. Cell reports 37: 109848. https://doi.org/10.1016/j.celrep.2021.109848.

Feasey NA, Dougan G, Kingsley RA, Heyderman RS, Gordon MA. 2012. Invasive non-typhoidal salmonella disease: an emerging and neglected tropical disease in Africa. *Lancet (London*, England) 379: 2489–2499. https://doi.org/10.1016/s0140-6736(11)61752-2.

Flannagan RS, Cosío G, Grinstein S. 2009. Antimicrobial mechanisms of phagocytes and bacterial evasion strategies. Nature reviews. Microbiology 7: 355–366. https://doi.org/10.1038/nrmicro2128.

Franklin RB, Costello LC. 2009. The important role of the apoptotic effects of zinc in the development of cancers. Journal of cellular biochemistry 106: 750–757. https://doi.org/10.1002/jcb.22049.

Gao H, Dai W, Zhao L, Min J, Wang F. 2018. The role of zinc and zinc homeostasis in macrophage function. Journal of immunology research 2018: 6872621. https://doi.org/10.1155/2018/6872621.

Gao H, Zhao L, Wang H, Xie E, Wang X, Wu Q, Yu Y, He X, Ji H, Rink L, Min J, Wang F. 2017. Metal transporter Slc39a10 regulates susceptibility to inflammatory stimuli by controlling macrophage survival. Proceedings of the National Academy of Sciences of the United States of America 114: 12940–12945. https://doi.org/10.1073/pnas.1708018114.

GBD. 2017. Estimates of global, regional, and national morbidity, mortality, and aetiologies of diarrhoeal diseases: a systematic analysis for the Global Burden of Disease Study 2015. Lancet Infect Dis 17: 909–948. https://doi.org/10.1016/s1473-3099(17)30276-1.

Golan Y, Berman B, Assaraf YG. 2015. Heterodimerization, altered subcellular localization, and function of multiple zinc transporters in viable cells using bimolecular fluorescence complementation. The Journal of biological chemistry 290: 9050–9063. https://doi.org/10.1074/jbc.M114.617332.

Gordon S, Taylor PR. 2005. Monocyte and macrophage heterogeneity. Nature reviews. Immunology 5: 953–964. https://doi.org/10.1038/nri1733.

Green LC, Wagner DA, Glogowski J, Skipper PL, Wishnok JS, Tannenbaum SR. 1982. Analysis of nitrate, nitrite, and [15N]nitrate in biological fluids. Analytical biochemistry 126: 131–138. https://doi.org/10.1016/0003-2697(82)90118-x.

Guo L, Lichten LA, Ryu MS, Liuzzi JP, Wang F, Cousins RJ. 2010. STAT5-glucocorticoid receptor interaction and MTF-1 regulate the expression of ZnT2 (Slc30a2) in pancreatic acinar cells. Proceedings of the National Academy of Sciences of the United States of America 107: 2818–2823. https://doi.org/10.1073/pnas.0914941107.

Haase H, Ober-Blöbaum JL, Engelhardt G, Hebel S, Heit A, Heine H, Rink L. 2008. Zinc signals are essential for lipopolysaccharide-induced signal transduction in monocytes. Journal of immunology (Baltimore, Md. : 1950) 181: 6491-6502. https://doi.org/10.4049/jimmunol.181.9.6491.

He X, Ge C, Xia J, Xia Z, Zhao L, Huang S, Wang R, Pan J, Cheng T, Xu PF, Wang F, Min J. 2023. The zinc transporter SLC39A10 plays an essential role in embryonic hematopoiesis. *Advanced science (Weinheim, Baden-Wurttemberg*, Germany): e2205345. https://doi.org/10.1002/advs.202205345.

Helaine S, Thompson JA, Watson KG, Liu M, Boyle C, Holden DW. 2010. Dynamics of intracellular bacterial replication at the single cell level. Proceedings of the National Academy of Sciences of the United States of America 107: 3746–3751. https://doi.org/10.1073/pnas.1000041107.

Hernandez LD, Pypaert M, Flavell RA, Galán JE. 2003. A *Salmonella* protein causes macrophage cell death by inducing autophagy. J Cell Biol 163: 1123–1131. https://doi.org/10.1083/jcb.200309161.

Hersh D, Monack DM, Smith MR, Ghori N, Falkow S, Zychlinsky A. 1999. The *Salmonella* invasin SipB induces macrophage apoptosis by binding to caspase-1. Proceedings of the National Academy of Sciences of the United States of America 96: 2396–2401. https://doi.org/10.1073/pnas.96.5.2396.

Jeon KI, Jeong JY, Jue DM. 2000. Thiol-reactive metal compounds inhibit NF-kappa B activation by blocking I kappa B kinase. Journal of immunology (Baltimore, Md. : 1950) 164: 5981-5989. https://doi.org/10.4049/jimmunol.164.11.5981.

Kambe T, Tsuji T, Hashimoto A, Itsumura N. 2015. The physiological, biochemical, and molecular roles of zinc transporters in zinc homeostasis and metabolism. Physiological reviews 95: 749–784. https://doi.org/10.1152/physrev.00035.2014.

Kimura T, Kambe T. 2016. The functions of metallothionein and ZIP and ZnT transporters: an overview and perspective. International journal of molecular sciences 17: 336. https://doi.org/10.3390/ijms17030336.

Krężel A, Maret W. 2017. The functions of metamorphic metallothioneins in zinc and copper metabolism. International journal of molecular sciences 18. https://doi.org/10.3390/ijms18061237.

Kuźmicka W, Manda-Handzlik A, Cieloch A, Mroczek A, Demkow U, Wachowska M, Ciepiela O. 2020. Zinc Supplementation Modulates NETs Release and Neutrophils’ Degranulation. Nutrients 13. https://doi.org/10.3390/nu13010051.

Lee SR. 2018. Critical role of zinc as either an antioxidant or a prooxidant in cellular systems. Oxidative medicine and cellular longevity 2018: 9156285. https://doi.org/10.1155/2018/9156285.

Lewis SM, Williams A, Eisenbarth SC. 2019. Structure and function of the immune system in the spleen. Science immunology 4. https://doi.org/10.1126/sciimmunol.aau6085.

Lichten LA, Cousins RJ. 2009. Mammalian zinc transporters: nutritional and physiologic regulation. Annual review of nutrition 29: 153–176. https://doi.org/10.1146/annurev-nutr-033009-083312.

Liu MJ, Bao S, Gálvez-Peralta M, Pyle CJ, Rudawsky AC, Pavlovicz RE, Killilea DW, Li C, Nebert DW, Wewers MD, Knoell DL. 2013. ZIP8 regulates host defense through zinc-mediated inhibition of NF-κB. Cell reports 3: 386–400. https://doi.org/10.1016/j.celrep.2013.01.009.

Liuzzi JP, Cousins RJ. 2004. Mammalian zinc transporters. Annual review of nutrition 24: 151–172. https://doi.org/10.1146/annurev.nutr.24.012003.132402.

Malavolta M, Basso A, Piacenza F, Giacconi R, Costarelli L, Pierpaoli S, Mocchegiani E. 2012. Survival study of metallothionein-1 transgenic mice and respective controls (C57BL/6J): influence of a zinc-enriched environment. Rejuvenation research 15: 140–143. https://doi.org/10.1089/rej.2011.1261.

Mastroeni P, Grant A, Restif O, Maskell D. 2009. A dynamic view of the spread and intracellular distribution of Salmonella enterica. Nature reviews. Microbiology 7: 73–80. https://doi.org/10.1038/nrmicro2034.

Moskovskich A, Goldmann U, Kartnig F, Lindinger S, Konecka J, Fiume G, Girardi E, Superti-Furga G. 2019. The transporters SLC35A1 and SLC30A1 play opposite roles in cell survival upon VSV virus infection. Scientific reports 9: 10471. https://doi.org/10.1038/s41598-019-46952-9.

Nishito Y, Kambe T. 2019. Zinc transporter 1 (ZNT1) expression on the cell surface is elaborately controlled by cellular zinc levels. The Journal of biological chemistry 294: 15686–15697. https://doi.org/10.1074/jbc.RA119.010227.

Palmiter RD. 2004. Protection against zinc toxicity by metallothionein and zinc transporter 1. Proceedings of the National Academy of Sciences of the United States of America 101: 4918–4923. https://doi.org/10.1073/pnas.0401022101.

Pfaffl MW. 2001. A new mathematical model for relative quantification in real-time RT-PCR. Nucleic acids research 29: e45. https://doi.org/10.1093/nar/29.9.e45.

Pham THM, Brewer SM, Thurston T, Massis LM, Honeycutt J, Lugo K, Jacobson AR, Vilches-Moure JG, Hamblin M, Helaine S, Monack DM. 2020. *Salmonella*-driven polarization of granuloma macrophages antagonizes TNF-mediated pathogen restriction during persistent infection. Cell host & microbe 27: 54–67.e55. https://doi.org/10.1016/j.chom.2019.11.011.

Prasad AS, Bao B, Beck FW, Sarkar FH. 2011. Zinc-suppressed inflammatory cytokines by induction of A20-mediated inhibition of nuclear factor-κB. Nutrition (Burbank, Los Angeles County, Calif.) 27: 816–823. https://doi.org/10.1016/j.nut.2010.08.010.

Rossi DC, Figueroa JAL, Buesing WR, Candor K, Blancett LT, Evans HM, Lenchitz R, Crowther BL, 3rd, Elsegeiny W, Williamson PR, Rupp J, Deepe GS, Jr. 2021. A metabolic inhibitor arms macrophages to kill intracellular fungal pathogens by manipulating zinc homeostasis. The Journal of clinical investigation 131. https://doi.org/10.1172/jci147268.

Schlesinger L, Arevalo M, Arredondo S, Lönnerdal B, Stekel A. 1993. Zinc supplementation impairs monocyte function. Acta paediatrica (Oslo, Norway: 1992) 82: 734-738. https://doi.org/10.1111/j.1651-2227.1993.tb12548.x.

Shembade N, Ma A, Harhaj EW. 2010. Inhibition of NF-kappaB signaling by A20 through disruption of ubiquitin enzyme complexes. *Science (New York*, N.Y.) 327: 1135–1139. https://doi.org/10.1126/science.1182364.

Shusterman E, Beharier O, Levy S, Zarivach R, Etzion Y, Campbell CR, Lee IH, Dinudom A, Cook DI, Peretz A, Katz A, Gitler D, Moran A. 2017. Zinc transport and the inhibition of the L-type calcium channel are two separable functions of ZnT-1. Metallomics : integrated biometal science 9: 228–238. https://doi.org/10.1039/c6mt00296j.

Smith PJ, Wiltshire M, Furon E, Beattie JH, Errington RJ. 2008. Impact of overexpression of metallothionein-1 on cell cycle progression and zinc toxicity. American journal of physiology. Cell physiology 295: C1399–1408. https://doi.org/10.1152/ajpcell.00342.2008.

Stocks CJ, von Pein JB, Curson JEB, Rae J, Phan MD, Foo D, Bokil NJ, Kambe T, Peters KM, Parton RG, Schembri MA, Kapetanovic R, Sweet MJ. 2021. Frontline Science: LPS-inducible SLC30A1 drives human macrophage-mediated zinc toxicity against intracellular Escherichia coli. Journal of leukocyte biology 109: 287–297. https://doi.org/10.1002/jlb.2hi0420-160r.

Swirski FK, Nahrendorf M, Etzrodt M, Wildgruber M, Cortez-Retamozo V, Panizzi P, Figueiredo JL, Kohler RH, Chudnovskiy A, Waterman P, Aikawa E, Mempel TR, Libby P, Weissleder R, Pittet MJ. 2009. Identification of splenic reservoir monocytes and their deployment to inflammatory sites. *Science (New York*, N.Y.) 325: 612–616. https://doi.org/10.1126/science.1175202.

Wertz IE, O’Rourke KM, Zhou H, Eby M, Aravind L, Seshagiri S, Wu P, Wiesmann C, Baker R, Boone DL, Ma A, Koonin EV, Dixit VM. 2004. De-ubiquitination and ubiquitin ligase domains of A20 downregulate NF-kappaB signalling. Nature 430: 694–699. https://doi.org/10.1038/nature02794.

Wessels I, Fischer HJ, Rink L. 2021. Dietary and physiological effects of zinc on the immune system. Annual review of nutrition 41: 133–175. https://doi.org/10.1146/annurev-nutr-122019-120635.

Wu A, Tymoszuk P, Haschka D, Heeke S, Dichtl S, Petzer V, Seifert M, Hilbe R, Sopper S, Talasz H, Bumann D, Lass-Flörl C, Theurl I, Zhang K, Weiss G. 2017. *Salmonella* utilizes zinc to subvert antimicrobial host defense of macrophages via modulation of NF-κB signaling. Infection and immunity 85. https://doi.org/10.1128/iai.00418-17.

Wu Q, Shen Y, Tao Y, Wei J, Wang H, An P, Zhang Z, Gao H, Zhou T, Wang F, Min J. 2017. Hemojuvelin regulates the innate immune response to peritoneal bacterial infection in mice. Cell discovery 3: 17028. https://doi.org/10.1038/celldisc.2017.28.

Wynn TA, Chawla A, Pollard JW. 2013. Macrophage biology in development, homeostasis and disease. Nature 496: 445–455. https://doi.org/10.1038/nature12034.

Xie QW, Kashiwabara Y, Nathan C. 1994. Role of transcription factor NF-kappa B/Rel in induction of nitric oxide synthase. The Journal of biological chemistry 269: 4705–4708.

Zhou Z, Wang L, Song Z, Saari JT, McClain CJ, Kang YJ. 2004. Abrogation of nuclear factor-kappaB activation is involved in zinc inhibition of lipopolysaccharide-induced tumor necrosis factor-alpha production and liver injury. The American journal of pathology 164: 1547–1556. https://doi.org/10.1016/s0002-9440(10)63713-3.

